# Microtubule-associated ROP interactors both delimit and transduce ROP signaling and regulate microtubule dynamics

**DOI:** 10.1101/2021.01.08.425872

**Authors:** Gil Feiguelman, Xiankui Cui, Hasana Sternberg, Ying Fu, Shaul Yalovsky

**Author notes:** Corresponding author: Shaul Yalovsky, School of Plant Sciences and Food Security, Tel Aviv University, Tel Aviv 6997801, Israel.

## Abstract

Evidence suggests that ICR proteins function as adaptors that mediate ROP signaling. Here, we studied the functions of ICR2 and its homologs ICR5 and ICR3. We showed that ICR2 is a microtubule-associated protein that regulates microtubule dynamics. ICR2 can retrieve activated ROPs from the plasma membrane, and it is recruited to a subset of ROP domains. Secondary cell wall pits in the metaxylem of *icr2* and *icr5 Arabidopsis* single mutants and *icr2*/*icr5* double and *icr2*/*icr5*/*icr3* triple mutants were denser and larger than those in wild-type Col-0 seedlings, implicating these three ICRs in restriction of ROP function. The *icr2* but not the *icr5* mutants developed split root hairs further implicating ICR2 in restriction of ROP signaling. Taken together, our results show that ICR2, and likely also ICR5 and ICR3, have multiple functions as ROP effectors and as regulators of microtubule dynamics.

## Introduction

ROP (Rho Of Plants) are the plant-specific subfamily of RHO superfamily of small G proteins. ROPs function as plasma membrane-anchored molecular switches that cycle between active, GTP-bound and inactive, GDP-bound states (Feiguelman et al., 2018). ROP activity is spatiotemporally controlled by regulatory proteins in plasma membrane-associated microdomains that contain active ROP. Guanine nucleotide Exchange Factors (GEFs) are known to positively regulate the activation of ROPs by facilitating nucleotide exchange (Basu et al., 2008; Berken et al., 2005; Thomas and Berken, 2010). GTPase Activating Proteins (GAPs) down-regulate ROP activity, and ROPs are recycled by GDP Dissociation Inhibitors (GDIs) (Boulter and Garcia-Mata, 2010; DerMardirossian and Bokoch, 2005; Schaefer et al., 2011; Wu et al., 2000). In the active, GTP-bound state, ROPs interact with target effector proteins to perform their biological functions (Dvorsky and Ahmadian, 2004; Feiguelman et al., 2018). ROPs regulate a variety of cellular processes such as the organization and dynamics of the actin and microtubule cytoskeleton, endocytosis and exocytosis, and activation of NADPH oxidase and intracellular kinase cascades. ROP functions impact cell growth and shape, cytokinesis, subcellular protein localization, and responses to pathogens and abiotic stresses (Bloch and Yalovsky, 2013; Feiguelman et al., 2018; Kawano et al., 2014; Nagawa et al., 2010; Oda and Fukuda, 2014; Rivero et al., 2017; Yalovsky et al., 2008; Yang, 2008).

We have previously identified a family of ROP effectors that we designated “Interactors of Constitutively active ROP” (ICRs) (Lavy et al., 2007). The ICRs are coiled-coil domain-containing proteins that do not contain additional known structural or catalytic domains (Lavy et al., 2007). The ICRs do contain two conserved sequence motifs, an N-terminal QEEL and a C-terminal QWRKAA (Lavy et al., 2007). The ICRs are subdivided into two clades, which differ in molecular mass. In *Arabidopsis*, ICR1 (At1g17140, 38 kDa) and ICR4 (At1g78430, 36 kDA) represent the lower molecular weight clade, whereas ICR2 (At2g37080, 65 kDa), ICR3 (At5g60210, 63 kDa), and ICR5 (At3g53350, 45 kDa) represent the higher molecular weight clade.

Studies on ICR1 have shown that it is a microtubule-associated protein (MAP) that integrates auxin and Ca^2+^ signaling. It functions as a ROP-associated scaffold that interacts with specific group of proteins (Hazak et al., 2010; Hazak et al., 2019; Hazak et al., 2014; Lavy et al., 2007). ICR1 is recruited to the plasma membrane by ROPs and subsequently recruits the EF-hand calcium binding protein Calcium-dependent Modulator of ICR1 (CMI1) to cortical microtubules. This effect on CMI1 subcellular distribution influences its function (Hazak et al., 2019; Lavy et al., 2007).

Studies on ICR5 (also known as ROP Interacting Partner 3 (RIP3) and Microtubule Depletion Domain 1 (MIDD1)) revealed that it is a MAP that interacts with the microtubule-destabilizing kinesin, Kinesin13A (Mucha et al., 2010). Functional analysis of dedifferentiating tracheary elements showed that ICR5 regulates secondary cell wall deposition in differentiating metaxylem cells through an association with depolymerizing cortical microtubules in future secondary cell wall pits (Oda and Fukuda, 2012; Oda and Fukuda, 2013; Oda et al., 2010). It was proposed that ICR5 is recruited to plasma membrane domains by ROP11, where it promotes local microtubule breakdown, which in turn results in the formation of cell wall pits (Oda and Fukuda, 2012; Oda and Fukuda, 2013). A recent study showed that that ICR2 and ICR5 interact with the AGC1.5 protein kinase, which in turn phosphorylates ROPGEF4 and ROPGEF10 to promote root hair growth (Li et al., 2020).

Plant microtubules function dynamically to regulate cellular functions related to cell division, cell growth, cell shape formation, pathogen invasion, and abiotic stresses. Microtubule dynamics, including extension, shrinkage, catastrophe, and rescue, have been studied in plant cells (Ehrhardt and Shaw, 2006; Elliott and Shaw, 2018; Shaw et al., 2003). Although plant cells have unique plant microtubule structures and organization processes, microtubule dynamics are conserved in eukaryotes (Hamada, 2014a). Microtubules organize in several typical structures in the course of the cell cycle. During interphase, microtubules form cortical arrays beneath the plasma membrane. In contrast to animal and yeast cells, in plant cells, interphase microtubules organize without an organizing center and their plus and minus ends are distributed throughout the cell cortex (Ehrhardt, 2008; Ehrhardt and Shaw, 2006; Elliott and Shaw, 2018; Wasteneys and Ambrose, 2009; Yagi et al., 2018; Yi and Goshima, 2018).

The dynamic nature of the cortical microtubules and their ability to respond to diverse stimuli is governed by MAPs that regulate their nucleation, stability, crosslinking, severing, membrane interaction, and orientation (Hamada, 2014b). For example, movement of cellulose synthases (CesAs) in the membrane is driven by the synthesis of cellulose chains and overlays with cortical microtubules (Paredez et al., 2006). Cortical microtubules have been suggested to affect CesAs localization in the plasma membrane and to regulate their movement (Gutierrez et al., 2009). Several MAPs are known to mediate CesA and microtubule colocalization (Bringmann et al., 2012; Endler et al., 2015; Gu et al., 2010; Kesten et al., 2019; Lei et al., 2013; Li et al., 2012).

Though ICR proteins are MAPs (Hazak et al., 2019; Mucha et al., 2010; Oda and Fukuda, 2012; Oda and Fukuda, 2013; Oda et al., 2010) their effect on microtubule organization and how they affect ROP signaling are not well understood. In this work, we characterized the functions of ICR2 and ICR5 and analyzed the *icr2/icr5* double and *icr2/icr5/icr3* triple mutant phenotypes. Our results indicate that ICR2 function is associated with restriction of ROP signaling domains and that the function ICR5 in differentiating metaxylem cell differs from that previously proposed.

## Results

### ICR2 expression pattern

To analyze the expression pattern of ICR2 protein, the genomic sequence of *ICR2*, including 2225 bp of upstream promoter sequence, was fused to the sequence encoding the β-glucuronidase (GUS) reporter (*pICR2::ICR2_genomic_-GUS*). At 7 days after germination (DAG) ICR2 expression was observed near the root tip, specifically in the cell division zone, and in lateral root initials, lateral roots, vascular tissues, and root hairs (Figure 1A-D). In the hypocotyl and the cotyledons, ICR2-GUS expression was strong in vascular tissues, leaf primordia, and stomata linage cells (Figure 1E-F). Interestingly, although detected, the expression was lower in mature guard cells than in stomata linage cells (Figure 1G and H). In reproductive organs, ICR2-GUS was observed in developing floral tissue, the vasculature of pedicels and receptacles, sepals, the stamen filament, ovary and ovules, and developing seeds and siliques (Figure 1I-L).

**Figure 1:**
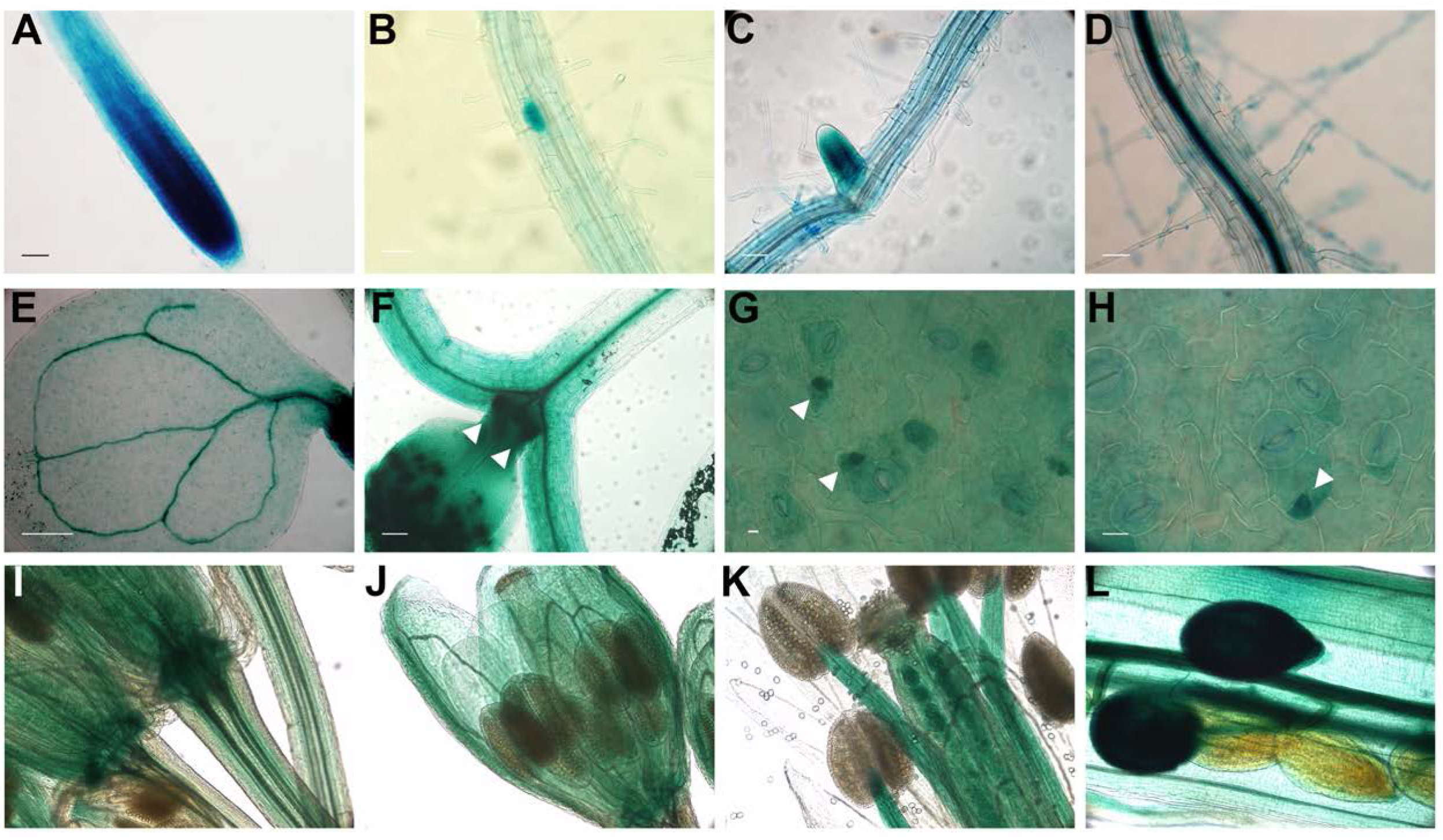
The expression pattern of ICR2. Expression of ICR2 was analyzed in *pICR2::ICR2-GUS* plants. Expression was detected in **(A)** the root tip, **(B)** lateral root initials, **(C)** developing lateral root, **(D)** stele and root hairs of root differentiation zone, **(E)** vascular tissues and developing stomata in cotyledons, **(F)** vasculature tissues and developing leaves hypocotyl (ICR2 is indicated by arrowheads), **(G and H)** meristemoids and developing guard cells (ICR2 is indicated by arrowheads), **(I)** vasculature in pedicels and receptacles, **(J)** vascular tissues in in sepals, **(K)** the stamen filaments, and **(L)** ovules, developing seeds, and siliques. Scale bars correspond to 50 μm for panels A-F and 20 μm for panels G-H. See also Figure 1-Supplement 1.

In agreement with the ICR2-GUS reporter data, gene expression data from the *Arabidopsis* eFP Browser (https://bar.utoronto.ca/efp/cgi-bin/efpWeb.cgi (Winter et al., 2007)) indicates that *ICR2* expression is higher in the shoot apex and in seeds than other tissues and higher during flower development than other stages (Figure 1-Supplement 1). A co-expression analysis using GENEVESTIGATOR (www.genevestigator.com/gv/ (Hruz et al., 2008)) indicated that the expression of *ICR2* is highly correlated with various MAPs and actin-associated proteins (Table S1). The co-expression data suggested that *ICR2* is involved in, or at the very least, up-regulated, during cell division, cytokinesis by cell plate formation, cell proliferation, regulation of the cell cycle, and cytoskeletal organization. Although many of the co-expressed genes are uncharacterized, the strong correlations of *ICR2* levels with levels of *ICR3* and *ICR4* suggest that they either function together or that there is some functional redundancy among these ICR family members. Interestingly, *MAP65-2*, which is the mRNA with expression most highly correlated with *ICR2* levels, is a coiled coil-containing microtubule-stabilizing protein involved in microtubule bundling in both interphase and cytokinetic microtubule arrays (Guo et al., 2009; Lucas et al., 2011; Lucas and Shaw, 2012). The GUS reporter expression data are in line with the transcriptomic data and indicate that *ICR2* is highly expressed in meristem and dividing cells, developing stomata, flower organs, ovules, and seeds. The relatively high expression detected in vascular tissues and root hairs suggests that ICR2 may function in these tissues and cells.

### Generation of single and double mutants *of ICR2* and *ICR5* and the *ICR2*, *ICR5*, and *ICR3* triple mutant

In order to characterize the function of ICR2, mutants were either obtained or generated. The *icr2-1* (*GK567F02*), *icr2-2* (*GK281B01*), and *icr2-3* (*GK159B08*) *T-DNA* mutants, which are part of the GABI KAT seed stock (Kleinboelting et al., 2012), were obtained from the European *Arabidopsis* Stock Centre (NASC) and are in the Columbia-0 (Col-0) background (Figure 2-Supplement 1). Further, multiplex genome editing by means of CRISPR/Cas9 (Bortesi and Fischer, 2015) was carried out in order to generate multiple mutant alleles in *ICR2*, *ICR3*, and *ICR5*. We identified two independent *icr5* single-mutant alleles, two independent *icr2/icr5* double-mutant alleles, and a single *icr2/icr3/ic*r5 triple mutant (Figure 2-Supplement 2). The CRISPR/Cas9-mediated genome editing generated InDels that resulted in mutant genes that encoded truncated proteins from which most residues were missing and were therefore very likely inactive. Hence all mutants were considered nulls.

### ICR2, ICR5, and ICR3 negatively regulate metaxylem pit formation

Previous work indicated that ICR5 is required for the formation of secondary cell wall pits in the metaxylem (MX) (Oda and Fukuda, 2012; Oda et al., 2010), but the phenotype of an *icr5* mutant has not been previously described. The creation of *icr2* and *icr5* single and double mutant plants, as well as the *icr2/icr3/icr5* triple mutant, enabled analysis of the functions of these ICRs in MX pit formation (Figure 2).

**Figure 2:**
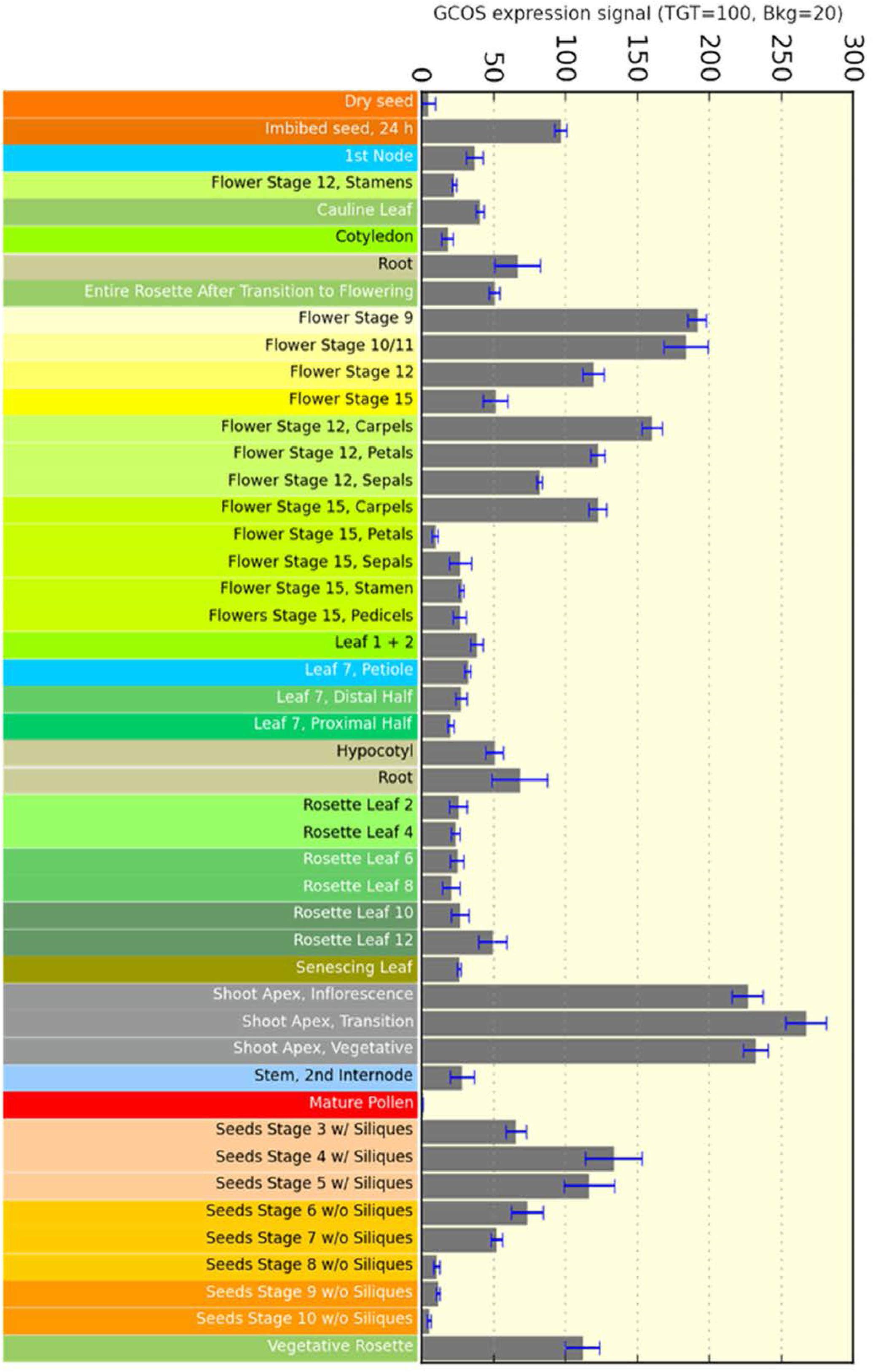
Metaxylem secondary cell wall pit size and density is regulated by ICRs. **(A-K)** DIC images of seedlings at 8 DAG, taken 3 or 4 cells shootward from the initiation of metaxylem differentiation. Scale bars, 10 μm. **(L)** Examples of pit area (yellow) and density (number of pits divided by area of red polygon) measurements. Scale bar, 10 μm. **(M)** Mean pit area (μm^2^). n>100 pits. (**N**) Pit density (number of pits per 1000 μm^2^). n>10 cells. Statistical analyses by one-way ANOVA. For panel M: F(10, 1400)=49.4836, p<0.0001 and for panel N: F(10, 122)=17.0351, p<0.0001. Means with different letters are significantly different (Tukey’s HSD, p<0.05). The boxes are the interquartile ranges, the whiskers represent the 1^st^ and 4^th^ quartiles, and the lines are the averages. See source data-Figure 2M, Figure 2N. See also Figure 2-Supplement 1, Figure 2-Supplement 2.

Surprisingly, contrary to previous predictions regarding the function of ICR5, the analysis of *icr2* and *icr5* single mutants revealed that they have significantly larger and denser pits than Col-0 plants. Hence pit formation was enhanced rather than suppressed in the *icr5*-null background. The size and the density of the pits in *icr2/icr5* and *icr2/icr3/icr5* double and triple mutants are increased compared to *icr2* and *icr5* single mutants, indicating that these three ICRs have redundant functions in the regulation of pit formation. Importantly, pit size and density were partially complemented in double transgenic plants expressing a genomic clone of ICR2 fused to three repeats of the YFP variant YPet, under regulation of the *ICR2* promoter, and ICR2 fused to the microtubule marker RFP-MBD (*icr2-1 X UBQ10::RFP-MBD X ICR2-3xYPet* and *icr2-2 X UBQ10::RFP-MBD X ICR2-3xYPet*).

### ICR5 but not ICR2 regulates protoxylem secondary cell wall deposition

To analyze whether ICR2, ICR3 and ICR5 have additional roles during vascular differentiation, specifically in secondary cell wall deposition, the density of developing protoxylem (PX) lignin coils was measured by imaging lignin auto-fluorescence (Figure 3A-J). The PX lignin coils in *icr2* mutants were similar to those of Col-0, whereas the *icr5* single mutants as well as the *icr2/icr5* and *icr2/icr3/icr5* mutants had denser lignin deposition than Col-0 (Figure 3K). This finding implicated ICR5 in regulation of secondary cell wall deposition in PX. As there was no additive phenotype in double and triple mutants, we reason that ICR2 and ICR3 do not function in the PX.

**Figure 3:**
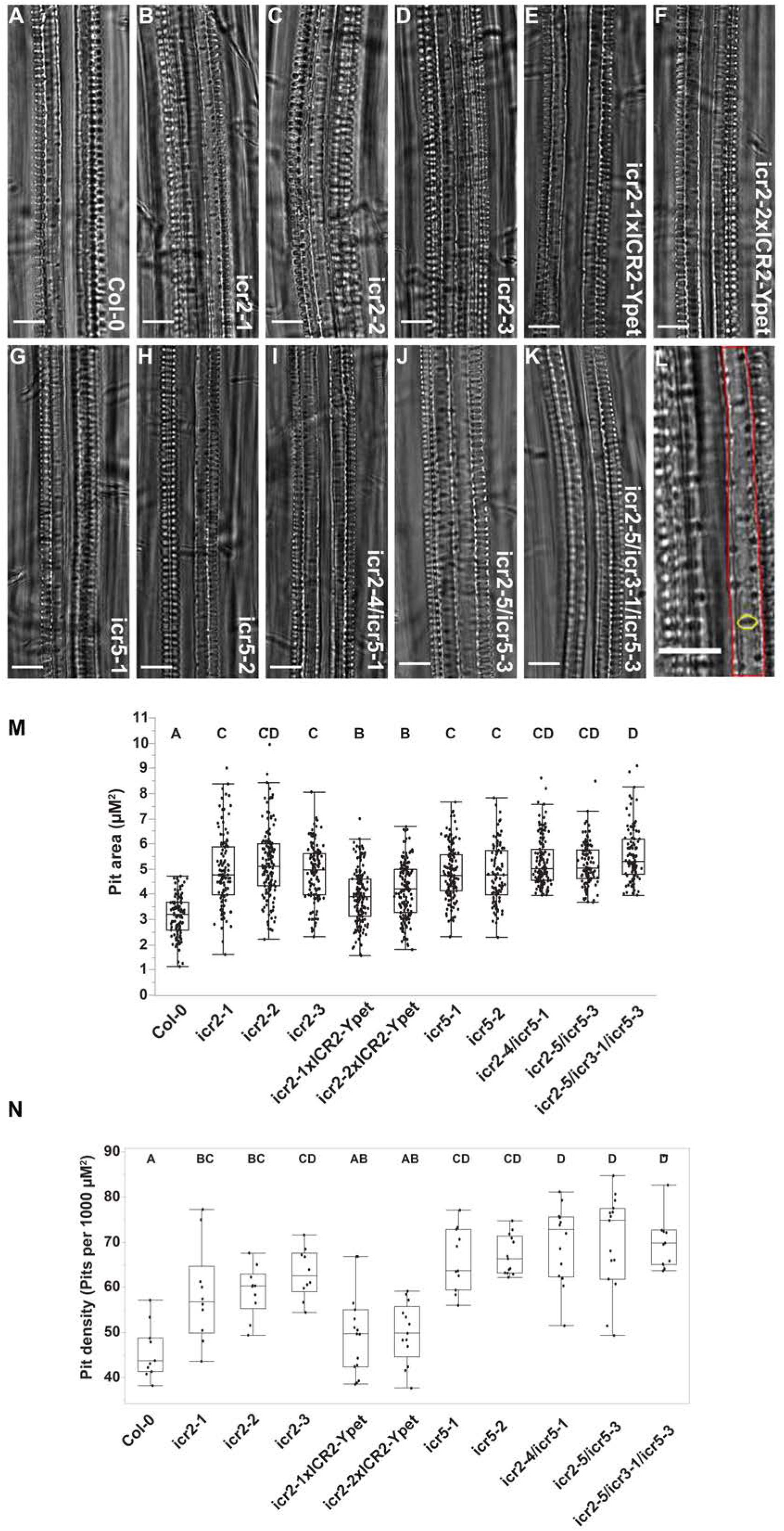
Protoxylem secondary cell wall coil density is regulated by ICR5. **(A-I)** Lignin auto-fluorescence in PX cells, imaged 3 or 4 cells shootward for initiation of differentiation. Scale bars, 20 μm. **J)** An exemplary image showing how the distance between the lignin coils was measured along white line. Images are Z-stacks of 8-11 confocal planes, 20x lens. **K)** Mean distance in μm between the lignin coils. Statistical analysis by one-way ANOVA: F(8, 86)=34.4972, p<0.0001. Means with different letters are significantly different (Tukey’s HSD, p<0.05). The boxes are the interquartile ranges, the whiskers represent the 1^st^ and 4^th^ quartiles, and the lines are the averages. n=10 for each line. See source data-Figure 3K.

### *icr2* mutants exhibit a deformed, branched root hair phenotype

ICR2 expression was detected in root hair (Figure 1D). Because ROP signaling plays central role in root hair tip growth (Bloch et al., 2005; Bloch et al., 2011; Carol et al., 2005; Chai et al., 2016; Denninger et al., 2019; Duan et al., 2010; Jones et al., 2002; Kang et al., 2017; Molendijk et al., 2001; Nakamura et al., 2018; Wan et al., 2017), we asked whether single, double, and triple *icr* mutants develop abnormal root hairs. All the plants with *ICR2* mutant alleles exhibited deformed, branched root hair phenotypes (Figure 4A-K). In contrast, *icr5* root hairs were normal, and the double and triple mutants showed no additive effects (Figure 4L). Further, there was partial complementation of the split root hair phenotype in *icr2-1* and *icr2-2* by ICR2-YPet (Figure 4L). These data indicate that the function of ICRs in root hair growth regulation is not redundant. To observe whether mutations in genes encoding ICRs have any other effect on root hair development, we measured the distance of the first root hair bulge from the root tip (Figure 4-Supplement 1A), the root hair length (Figure 4-Supplement 1B) and root hair density (Figure 4-Supplement 1C). None of these characteristics differed in the *icr* mutants compared to Col-0 plants. This finding suggested that ICR2 is involved specifically in polarity maintenance of growing root hairs but not in root hair initiation. This is the first evidence that a ROP interactor affects root hair polarity.

**Figure 4:**
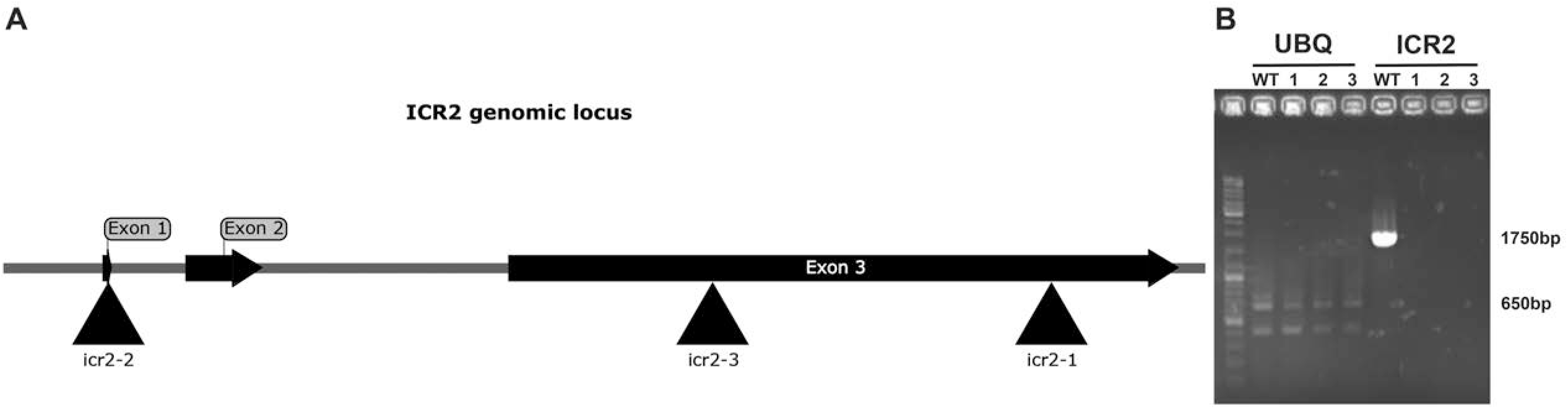
*icr2* mutants develop deformed split root hairs. **(A-K)** Representative images for each genotype. Arrowheads highlight split root hairs. Scale bars, 50 μm. **(L)** Percentage of abnormal root hairs in seedlings at 8 DAG. n=5 seedlings for each line. See source data-Figure 4L. See also Figure 4-Supplement 1.

### Interaction of ICR2 with microtubules and ROPs *in vivo*

Similar to secondary cell wall pits in MX, the split root hair phenotype has been associated with perturbation in microtubules and was described for several MAP mutants (Kang et al., 2017; Sakai et al., 2008; Whittington et al., 2001; Yang et al., 2007; Zhang et al., 2015). We discovered ICR2 in a yeast two-hybrid screen with constitutively active ROP10 (rop10^CA^) as bait (Lavy et al., 2007). Transient expression in *Nicotiana benthamiana* leaf epidermis cells showed that, when individually expressed, ICR2 localizes along cortical microtubules (Figure 5A, Figure 5-Supplement 1), whereas ROP11 localizes to the plasma membrane (Figure 5B) as has previously been demonstrated (Bloch et al., 2005; Lavy et al., 2002; Lavy and Yalovsky, 2006). The localization of ICR2 on microtubules was verified by treatment with the microtubule inhibitor oryzalin, which resulted in the disappearance of ICR2-labeled microtubules and a shift of ICR2 to the cytoplasm (Figure 5-Supplement 1B). Localization of ICR2 to microtubules was further verified by co-expression with the microtubule marker RFP-MBD with and without oryzalin treatment (Figure 5-Supplement 1F-H).

**Figure 5:**
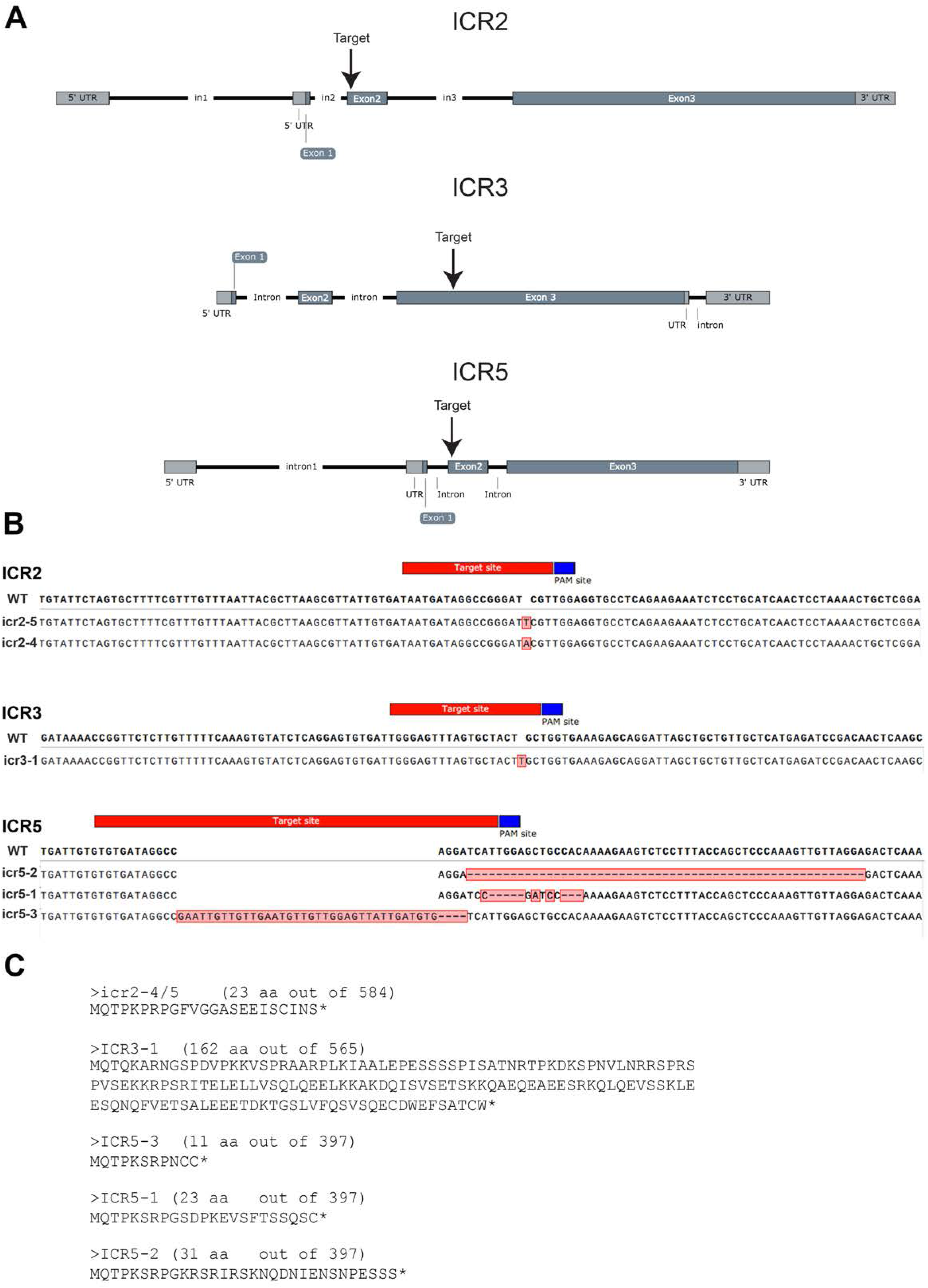
ICR2 is a microtubule-associated protein that interacts with ROP GTPases. **(A)** Image of *N. benthamiana* leaf epidermis that transiently express ICR2-3xYPet. Scale bar, 10 μm. **(B)** Image of *N. benthamiana* leaf epidermis that transiently express GFP-ROP11. Scale bar, 10 μm. **(C-L)** BiFC images of *N. benthamiana* leaf epidermis that transiently express ICR2-YC and **C)** YN-rop6^CA^, **D)** YN-rop9^CA^, **E)** YN-rop10^CA^, **F)** YN-rop11^CA^, **G)** YN-ROP2, **H)** YN-ROP4, **I)** YN-ROP6, **J)** YN-ROP9, **K)** YN-ROP10, and **L)** YN-ROP11. Scale bars for panels C-F, 10 μm and for panels G-L, 20 μm. YFP signal for panels G-I was separated by linear unmixing. Images for panels A, C-F are Z-projections of multiple confocal sections. **(M)** Yeast two-hybrid assays of ICR2 with ROP2, ROP4, ROP6, ROP9, ROP10, and ROP11. -LT, Leu-, Trp-deficient medium; -LTH, Leu-, Trp-, His-deficient medium; 3AT – 3-amino-1,2,4-triazole. Numbers above the panels denote dilution series. See also Figure 5-Supplement 1.

The interaction of ICR2 with ROPs was examined using bimolecular fluorescence complementation (BiFC). ICR2 fused at its carboxy terminus to the C-terminal half of YFP (ICR2-YC) was transiently expressed in *N. benthamiana* leaf epidermis cells along with various ROPs fused at their N-termini to the N-terminal half of YFP (YN-ROPs). ICR2 interacted with constitutively active versions of the type-I ROP6 or type-II ROP9, ROP10, or ROP11 or wild-type ROP2, ROP4, or ROP6 (type-I ROPs) or wild-type ROP9, ROP10, or ROP11 (type-II ROPs). The complexes localized along cortical microtubules (Figure 5C-L). Furthermore, in yeast two-hybrid assays, ICR2 interacted with both type-I ROP2, ROP4, and ROP6 and with type-II ROP9, ROP10, and ROP11 (Figure 5M). Taken together, these results suggest that ICR2 is a microtubule-associated ROP interactor that may recruit ROPs to microtubules from the plasma membrane.

To further explore the interaction between different ROPs and ICR2, GFP-tagged ROP2, ROP4, ROP6, ROP9, ROP10, and ROP11 were co-expressed in *N. benthamiana* leaf epidermis with the catalytic Plant-specific ROP Nucleotide Exchanger (PRONE) domain of ROPGEF3 (GEF3p), GAP1, and mCherry-tagged ICR2 (ICR2mCh). No changes in the organization of ICR2mCh-labeled microtubules were found around ROP2 domains (Figure 6A), and faint ICR2mCh labeling was detected around the ROP4 domains (Figure 6B). However, significant clustering of ICR2mCh-tagged microtubules was found around domains containing ROP6, ROP9, and ROP10 (Figure 6C-E) and weaker but clearly visible microtubule reorganization was found around ROP11 domains (Figure 6F). These results suggest that ROPs may differ in their abilities to recruit ICR2-associated microtubules to specific domains in the plasma membrane. We hypothesize that ICR2 has a dual role: On one hand, it restricts ROP activity to domains in the plasma membrane, and on the other hand, it functions as a scaffold for ROP interactions with microtubules and possibly with other proteins.

**Figure 6:**
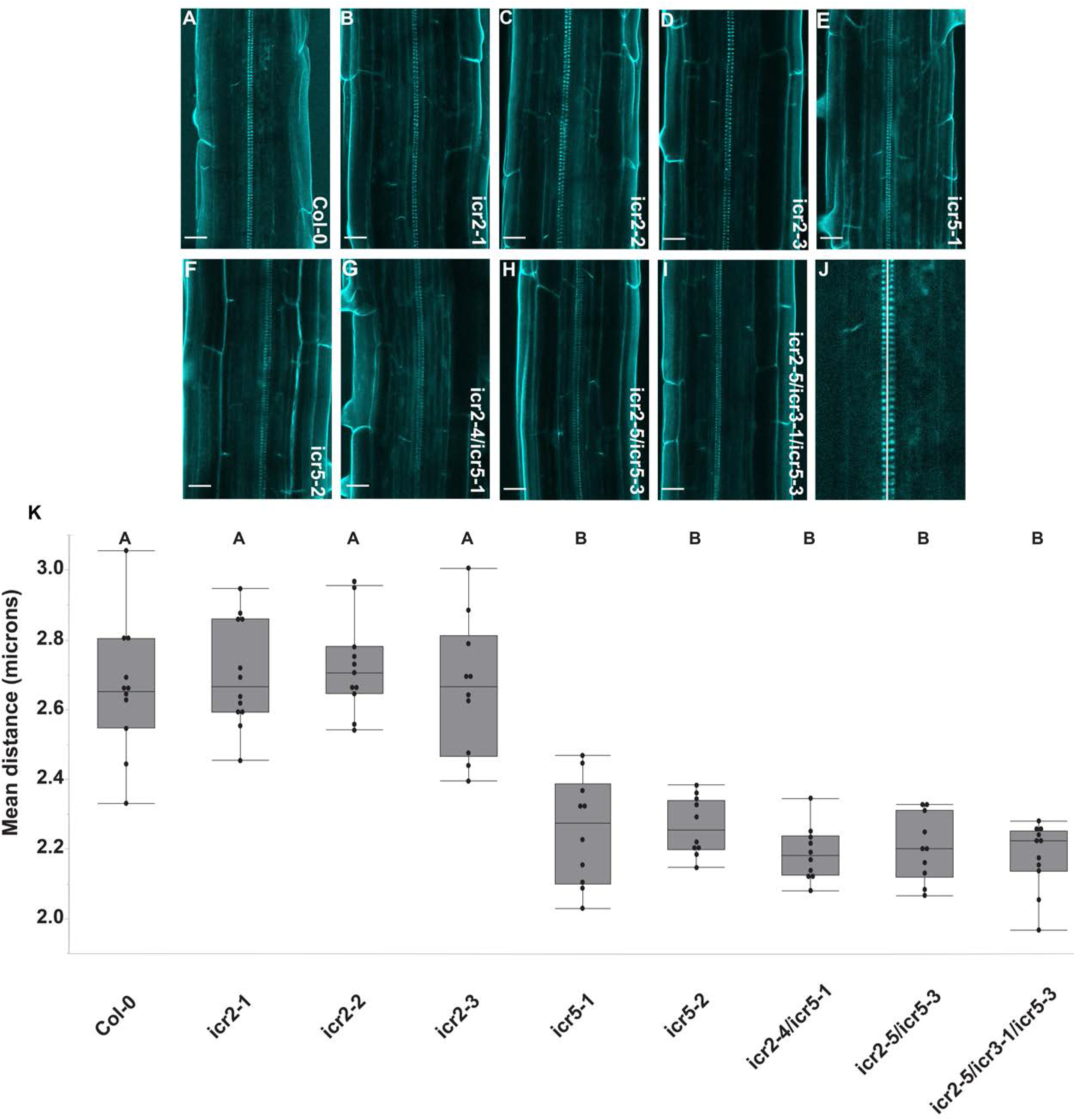
Microtubule-associated ICR2 is recruited to GEF3_PRONE_-GAP1-ROP domains. Images of *N. benthamiana* leaf epidermis that transiently express GFP-tagged ROPs (green) **(A)** ROP2, **(B)** ROP4, **(C)** ROP6, **(D)** ROP9, **(E)** ROP10, and **(F)** ROP11 with ICR2-mCherry (red), untagged GAP1, and the GEF3 PRONE domain (GEF3p). Scale bars, 10 μm in all panels.

### ICR1 and ICR2 bind microtubules *in vitro*

Both ICR1 and ICR2 localize to microtubules *in vivo* (Hazak et al., 2019; Mucha et al., 2010; Oda and Fukuda, 2012; Oda and Fukuda, 2013; Oda et al., 2010), but it is possible that their localization could have resulted from interaction with a third component rather than direct interaction with microtubules. To examine whether ICR1 and ICR2 are indeed MAPs, we tested their interactions with microtubules *in vitro* using three independent assays.

*Escherichia coli*-expressed, affinity-purified ICR1-His_6_ and ICR2-His_6_ at concentrations ranging between 1 to 10 μM were incubated with preformed taxol-stabilized microtubules. The protein mixtures were precipitated by centrifugation at 100,000 x *g*, and the precipitated proteins were separated by SDS-PAGE and visualized by Coomassie blue staining (Figure 7A, 7-Supplement 1A). The levels of precipitated ICR1-His_6_ and ICR2-His_6_ were quantified by densitometry of the relevant bands (Figure 7B, 7-Supplement 1B). MAP65, a known microtubule-interacting protein, was used a positive control. The binding of recombinant ICRs to microtubules was saturated at stoichiometries of 0.4 mol ICR1-His_6_ per mol of tubulin and 0.85 mol ICR2-His_6_ per mol of tubulin. This is in agreement with our findings that both ICR1 and ICR2 interact directly with microtubules and that the binding of ICR2 with microtubules is stronger than that of ICR1.

**Figure 7:**
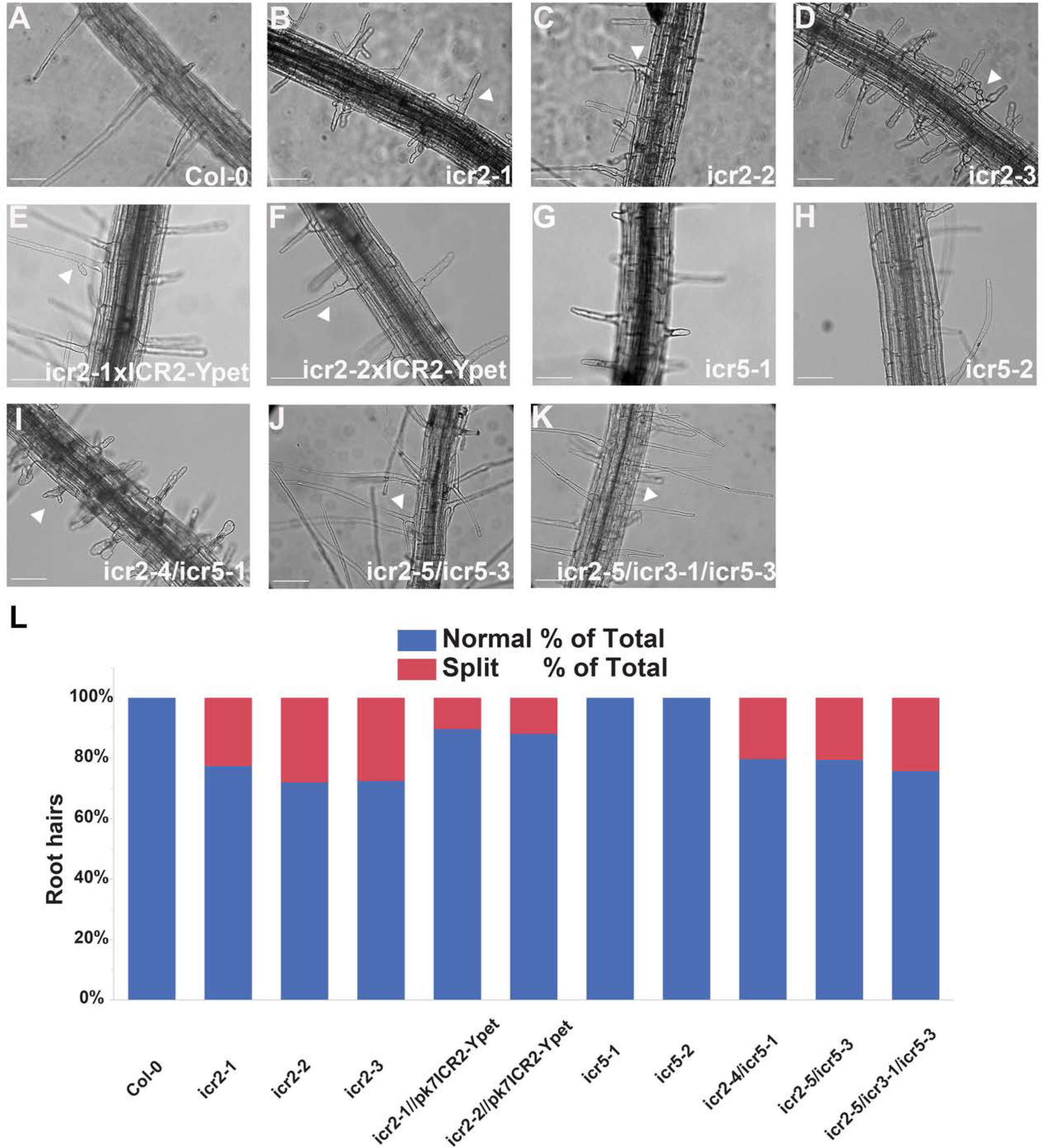
ICR2 interacts with microtubules *in vitro*. **(A)** Coomassie brilliant blue stained SDS-PAGE of recombinant ICR2-His_6_ co-sedimented with taxol-stabilized microtubules pre-polymerized from 5 μM tubulin. His-AtMAP65-1 was used as a positive control and GFP-His as a negative control. **(B)** Quantification of the ICR2-His_6_ band in panel A. The plot averages three replicates. **(C-H)** Immunofluorescence images of rhodamine-labeled tubulin (red) mixed with non-labeled tubulin and polymerized into microtubules in the presence of fluorescein-labeled ICR2-His_6_ (green). Arrowhead in panel D indicates ICR2 on a microtubule. Denatured ICR2-His was used as control. Scale bars, 10 μm. **(I-K)** Images of rhodamine-labeled tubulin bundling in the presence of 0, 0.1, or 2 μM ICR2. See source data-Figure 7B. See also Figure 7-Supplement 1, Figure 7-Supplement 2, Figure 7-Supplement 3.

Second, *in vitro* immuno-fluorescence assays were used to examine whether ICR1 and ICR2 colocalize with polymerized microtubules. To visualize microtubules, rhodamine-labeled tubulin was mixed with non-labeled tubulin and polymerized into microtubules in the presence of ICR1-His_6_ or ICR2-His_6_. Visualized by *in vitro* immuno-localization established that ICR1 and ICR2 are MAPs (Figure 7C-E, 7-Supplement 2A-C). Incubation with denatured ICR1-His_6_/ICR2-His_6_ were used as negative controls (Figure 7F-H, 7-Supplement 2D-F). ICR1 was distributed in individual punctae, whereas ICR2 was more evenly distributed along microtubule filaments suggesting stronger binding, in line with results of co-precipitation assays.

Third, we carried out an *in vitro* microtubule bundling assay. To this end, rhodamine-labeled tubulin was polymerized into microtubules in the presence of increasing concentrations of ICR1 or ICR2. With ICR1, microtubule bundling was detected only at the high concentration of 5 μM, whereas ICR2 caused bundling even at 0.1 μM (Figure 7I-K, 7-Supplement 3). Taken together, the bundling assays further confirmed that that ICR1 and ICR2 are MAPs and that binding of ICR1 to microtubules is weaker than ICR2 binding.

### ICR2 co-localizes with microtubules in all stages of the cell cycle

To characterize the subcellular localization of ICR2, we generated a marker composed of the genomic sequence of *ICR2* (including introns) with its promoter fused with the sequence for 3xYPet. To reduce potential steric hindrance a 33 amino acid linker was placed between ICR2 and the 3xYPet tag. To avoid potential mis-localization due to overexpression, the *pICR2::ICR2_genomic_:3xYpet* construct was transformed into two *icr2 T-DNA* insertion mutants, *icr2-1* and *icr2-2*, that also express the microtubule marker RFP-MBD (*icr2-1 X UBQ10::RFP-MBD* and *icr2-2 X UBQ10::RFP-MBD)*. Importantly, the *pICR2::ICR2_genomic_:3xYPet* fusion complemented the *icr2-1* and *icr2-2* pit formation and root hairs phenotypes, confirming its functionality (Figure 2-4). In the lateral root cap, root hairs, and in root epidermis cells, the ICR2:3xYPet fusion protein was observed on cortical microtubules (Figure 8A-I). The localization of ICR2 on microtubules was confirmed by colocalization with the microtubule marker RFP-MBD (Figure 8C, F, I, and J). The localization of ICR2 on microtubules in root hairs suggested that the split root hair phenotype of the *icr2* mutants is associated with ICR2 function on microtubules. Given that ICR2 can retrieve ROP2 from the plasma membrane to microtubules (Figure 5G) and given the similarities between the phenotypes of the *icr2* null and ROP2 gain-of-function mutants (Jones et al., 2002; Kang et al., 2017), ICR2 may restrict ROP2 signaling by recruiting it to microtubules.

**Figure 8:**
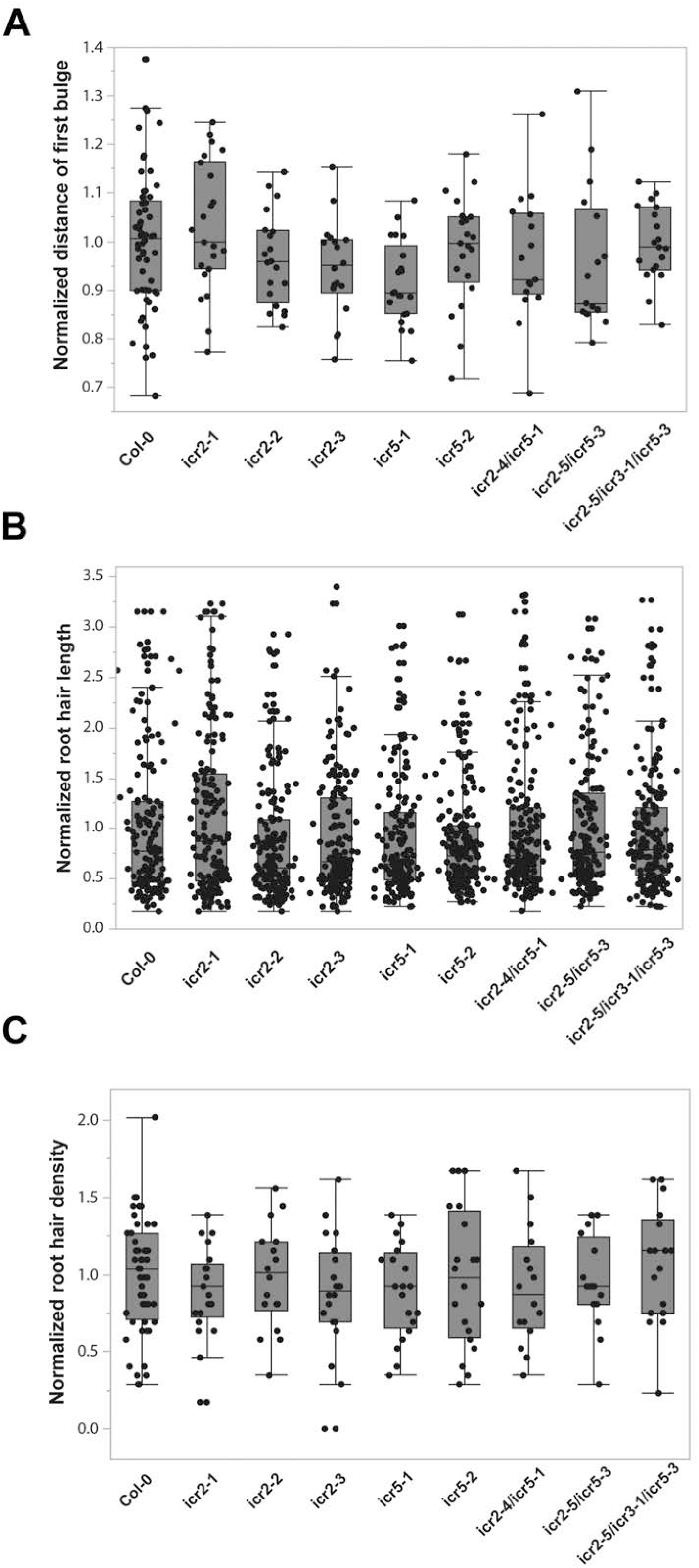
ICR2-3xYPet colocalizes with microtubules. **(A-C)** Images of *icr2-2* roots that express ICR2-3xYPet (green) and RFP-MBD (magenta). ICR2 was detected during interphase at the root tip, in the lateral root cap, and in dividing cells in the root cortex, as indicated by arrowheads. Scale bars, 10 μm. **(D-F)** Images of root hair shank in *icr2-2* plants that express ICR2-3xYPet (green) and RFP-MBD (magenta). Scale bars, 10 μm. **(G-I)** Images of differentiation/elongation zone epidermis in *icr2-2* plants that express ICR2-3xYPet (green) and RFP-MBD (magenta). Scale bars, 10 μm. **(J)** Fluorescence intensity profile of MRF-MBD and ICR2-3xYPet signals along the white lines in panels G-I. See source data-Figure 8J.

In dividing cells, ICR2 was colocalized with microtubules during mitosis (Figure 9A). ICR2-3xYPet colocalized with RFP-MBD in all mitotic stages including the preprophase band, the spindle during metaphase and anaphase, and the expanding phragmoplast microtubules in telophase (Figure 9B). Interestingly, the localization of ICR2 on microtubules during cell division coincided with the co-expression data, which showed high correlations with cell division cytoskeleton genes (Table S1).

**Figure 9:**
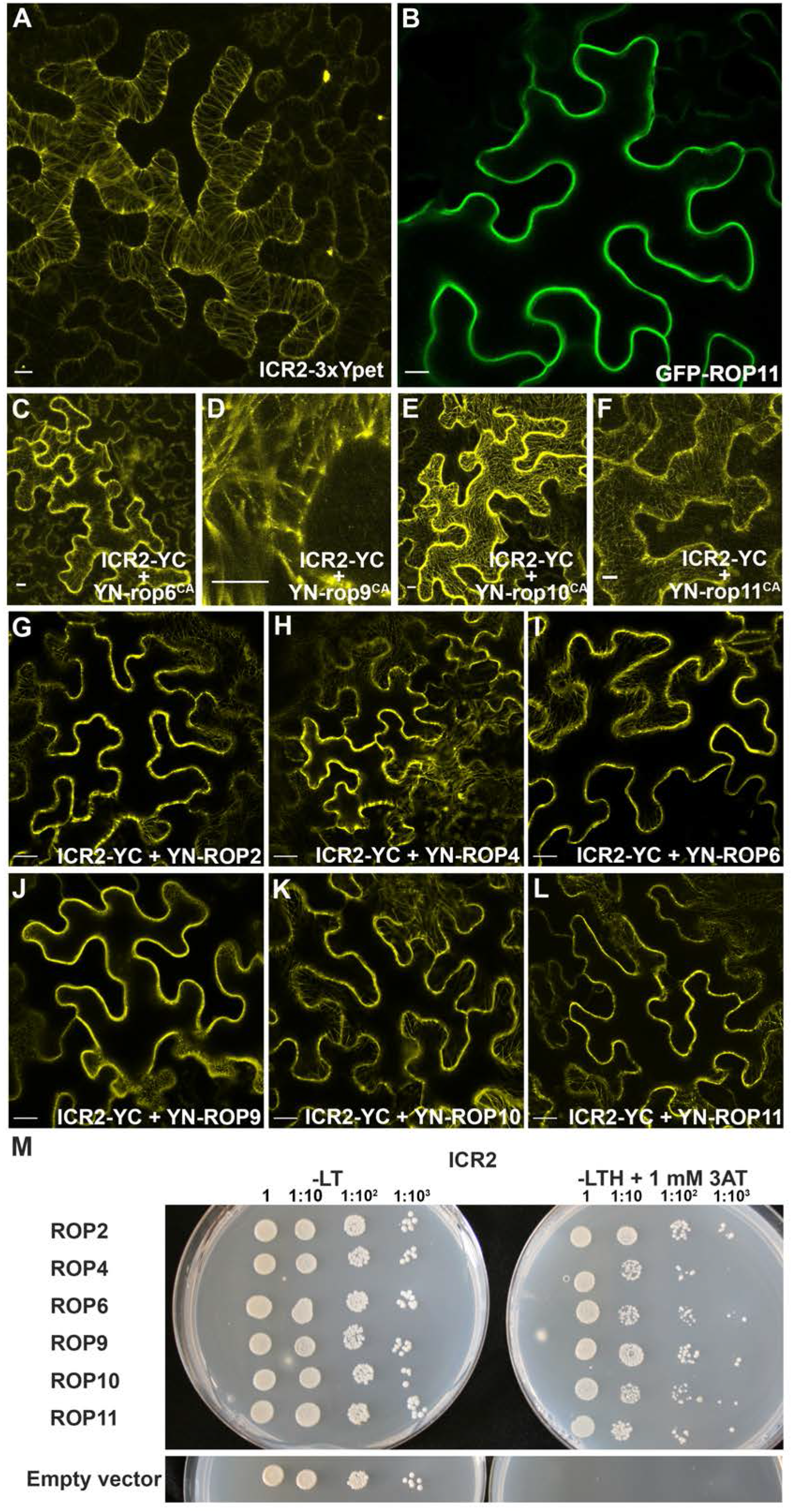
ICR2-3xYPet co-localizes with microtubules during all stages of cell division. **)A)** Image of lateral root cap of *icr2-2* plant expressing ICR2-3xYPet (green) and RFP-MBD (magenta). The expanding phragmoplast is indicated by arrowheads. Scale bars, 5 μm. **(B)** Tracking of a single cell undergoing cell division and cytokinesis in *icr2-2* plant expressing ICR2-3xYPet (green) and RFP-MBD (magenta). Colocalization is observed in all mitotic stages: preprophase band in prophase, spindle during metaphase and anaphase, and phragmoplast in telophase. Scale bars, 5 μm.

The pit phenotype of *icr2* mutants prompted us to examine ICR2 localization in vascular tissues. Unfortunately, with the experimental setup available to us for imaging of microtubules in the vasculature and the low expression levels of ICR2 made direct imaging impossible. To overcome this difficulty, we used bikinin-induced xylem/phloem dedifferentiation of leaf/cotyledon mesophyll cells (Kondo et al., 2016). The dedifferentiation of the mesophyll cells to cells harboring secondary cell wall occurred within 4 days and was visible using lignin auto fluorescence (Figure 10A-D). ICR2 expression was detected in differentiating xylem cells during the beginning of secondary cell wall deposition in cells that were still had chloroplasts (Figure 10A-D). ICR2-3xYPet fluorescence was not detected in fully differentiated cells (Figure 10A-D). Hence, ICR2 was expressed during relatively short time window at the onset of dedifferentiation indicating that its likely has specific function during xylem differentiation. The short time window in which ICR2 was expressed may also explain why it was difficult to detect it in root vascular tissues.

**Figure 10:**
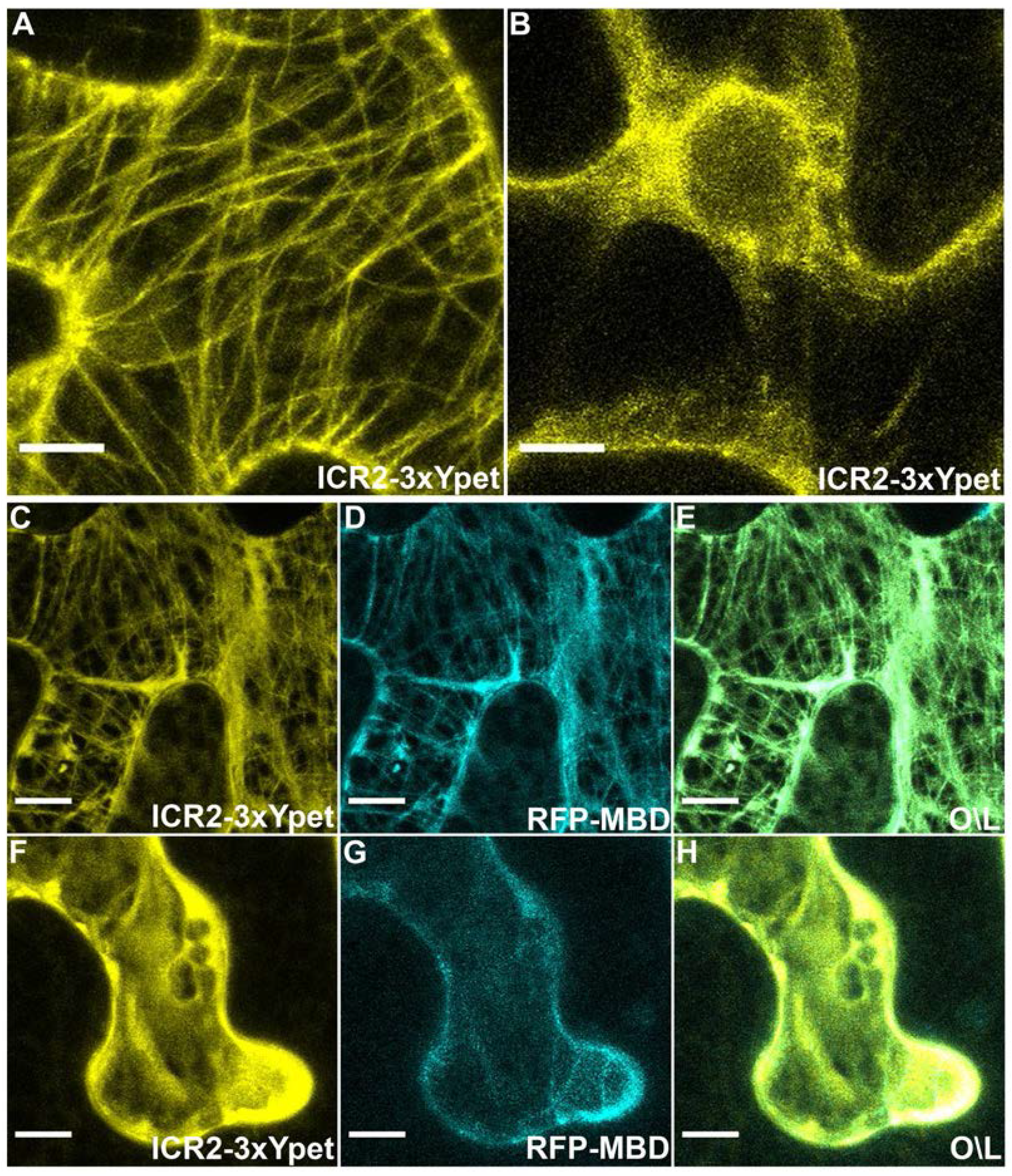
ICR2 is detected in differentiating xylem cells but not differentiated cells. **(A-D)** Images of secondary cell walls of cotyledon mesophyll cells undergoing dedifferentiation to tracheary elements: **A)** lignin autofluorescence (cyan), **B)** ICR2-3xYPet (yellow), **C)** chlorophyll (red), and **D)** overlay. Newly differentiating cells are indicated by arrows and fully differentiated cells by asterisks). Scale bars, 10 μm. **(E)** Fluorescence intensity profile of ICR2-3xYPet (yellow) lignin autofluorescence (cyan) along the line in panels A and B. See source data-Figure 10E.

In the dedifferentiating xylem cells, ICR2-3xYPet colocalized with lignified secondary cell wall (Figure 10E), whereas no fluorescence was detected in the pitted areas. Because the ICR2-3xYPet fluorescence was detected only in few cells and was absent in most of the cells that had secondary cell walls, it could not be a results of fluorescence channel spillover. Furthermore, the images were generated using spectral separation to ensure the presence of the YPet fluorescence. The strict colocalization of ICR2 with the secondary cell walls strongly suggests that it was localized along cortical microtubules. The pit phenotype of the *icr2* mutant implicates ICR2 in restriction of pit formation and regulation of pit size. In the developing MX, ICR2 may function by recruiting ROPs from the plasma membrane to microtubules.

### *icr2* mutants display altered microtubule organization and dynamics

The localization of ICR2 to microtubules at all stages of the cell cycle suggested that it may affect the organization and dynamics of microtubules. To test this, *icr2-1* and *icr2-2* plants were crossed with *UBQ10::RFP-MBD*, and analysis of microtubule dynamics was carried out on non-segregating double homozygous plants using high-frequency time-lapse imaging and tracking of individual microtubule filaments. The tracking data (Figure 11A) was used to create kymographs (Figure 11B), which were then used to calculate microtubule growth and shrinkage rates, the time spent in each condition, transition times, and pauses in growth/shrinkage. In root epidermal cells as well as root hairs, microtubule growth rates were significantly slower in the *icr2* mutants than Col-0 plants (p≤0.001) (Figure 11C). In contrast, shrinkage rates were lower only in the epidermis (Figure 11C). Additionally, time spent at pause was higher in mutant root epidermal cells than Col-0 plants (Figure 11-Supplement 1), and the transitions between filament growth, shrinkage, and pause occurred at higher frequency in the *icr2* mutants than in Col-0 plants (Figure 11-Supplement 2).

**Figure 11:**
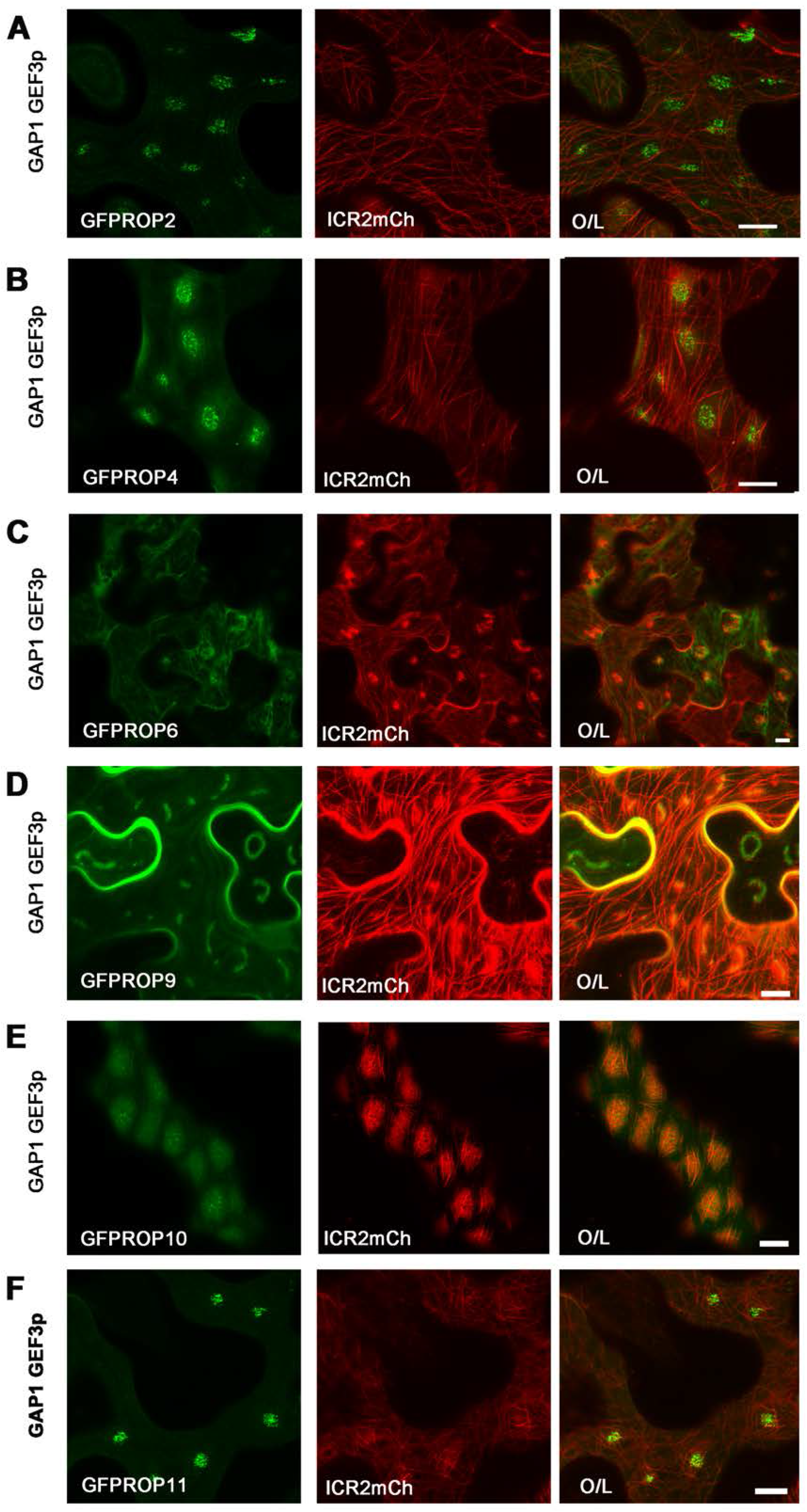
ICR2 affects microtubule dynamics in the root epidermis and root hairs. **(A)** Representative time-lapse imaging of RFP-MBD-labeled microtubules. Filament extension, pause, and shrinkage are labeled with green, blue, and red dots, respectively. Scale bars, 5 μm. **B)** Kymographs tracking RFP-MBD-labeled microtubule tips in *Col-0, icr2-1, icr2-2* seedlings. (**C)** Quantification of microtubule extension and shrinkage rates. Means with different letters are significantly different (Tukey’s HSD, p<0.05). One-way ANOVA results are in Table S3. The boxes are the interquartile ranges, the whiskers represent the 1^st^ and 4^th^ quartiles, and the lines are the averages. n>77 filaments per genotype. See source data-Figure 11. See also Figure 11-Supplement 1, Figure 11-Supplement-2.

Taken together the analysis of microtubule dynamics indicated that ICR2 regulates microtubule stability differently in different cell types. This cell-specific regulation of microtubule dynamics by ICR2 is consistent with the developmental phenotypes in root hair growth in *icr2* mutants but not the *icr5* single mutant (Figure 4). The secondary cell wall deposition phenotype of *icr5* but not *icr2* in PX (Figure 3) further supports cell- or tissue-specific function of ICR2. The MX pit patterning phenotype (Figure 2) suggests that ICR5, like ICR2, functions in a cell-specific manner.

## Discussion

### The functions of ICR2, ICR5, and ICR3

ICR proteins have been suggested to function as adaptors that mediate the interaction of ROPs with distinct target proteins (Lavy et al., 2007; Li et al., 2020; Mucha et al., 2010; Oda and Fukuda, 2012). The phenotypic analysis of the mutants presented in this work indicate that the function of the ICRs is more complex than previously thought. The mutant phenotypes indicate that at least ICR2, ICR3, and ICR5 also function as regulators that restrict ROP signaling, likely by recruiting ROPs from the plasma membrane to microtubules. Furthermore, the phenotypic analysis showed that the function of ICR2 and ICR5 is partially cell specific and that at least ICR2 may have ROP-independent functions in the regulation of microtubule dynamics. The data also suggest that ICR2 functions as effector for some ROPs. By recreating the naturally occurring active ROP domains in the plasma membrane, we were able to visualize how a subset of plasma-membrane anchored, active ROPs recruit the microtubule-associated ICR2 to these domains, thus relaying spatiotemporally regulated signal transduction into the cell.

### ICR2 is a MAP

The *in vitro* assays confirmed the ICR2 is a MAP. Our *in vitro* system also enabled us to compare the microtubule binding strength of ICR1 and ICR2. The co-sedimentation with microtubules revealed that ICR2 has a higher affinity for microtubules than ICR1. Whereas the binding of recombinant ICR2-His_6_ to microtubules was not saturated until 0.85 mol per mol of tubulin, saturation of ICR1-His_6_ was reached at only 0.35 mol per mol of tubulin. Similarly, the bundling assays showed that ICR1 caused microtubule bundling at 5 μM, whereas ICR2 induced microtubule bundling at the much lower concentration of 0.1 μM. The tighter binding of ICR2 to microtubules was also detected in the *in vitro* colocalization assays, which revealed an even distribution of ICR2 along microtubules at 0.5 μM, whereas ICR1 was detected as discrete punctae at 1 μM. It will be interesting to examine whether and how the differential binding of ICR1 and ICR2 to microtubules affects their functions as MAPs and as ROP signaling effectors.

### The involvement of ICR2, ICR5, and ICR3 in secondary cell wall patterning

ROP11 was previously implicated in regulation of secondary cell wall pits. A model involving ICR5 in the process was suggested previously (Oda and Fukuda, 2012; Oda et al., 2010). In this model, locally activated ROP11 recruits ICR5, leading to depolymerization of cortical microtubules in the future pit regions. Here we found that *icr2* and *icr5* single mutants have larger, denser pits compared to *Col-0*, whereas the *icr2/icr5* double and the *icr2/icr3/icr5* triple mutants have even larger pits and higher pit densities than the single mutants. These data indicate that ICR2, ICR3, and ICR5 have redundant functions in the regulation of pit formation in the MX. Additionally, expression of *ICR2-3xYPet* restored pit density in *icr2-1* and *icr2-2* backgrounds further confirming the function of ICR2 in regulation of pit formation. In dedifferentiating vascular cells ICR2 colocalized with secondary cell wall, indicating that its function is associated with the microtubules and secondary cell wall deposition rather than microtubule destabilization.

Furthermore, the microtubule dynamics analysis showed decreased microtubule growth rates in the *icr2* mutant background. If the primary function of ICR5 is induction of local microtubule destabilization through the recruitment of Kinesin13A, we would have expected that the size and number of pits would be reduced and not increased as seen in the *icr5* mutants. Thus, our data indicate that the functions of ICR5 as well as ICR2 and ICR3 are more complex than previously proposed. Possibly, ICR2, ICR3, and ICR5 restrict ROP domain activity by moving ROPs from the plasma membrane to microtubules. The higher density of secondary cell wall coils in the PX of *icr5* is in line with a role of ICR5 in destabilization of microtubules. Based on a combination of experimental work and computer simulation, Schneider et al. recently proposed that microtubule destabilization takes place during secondary cell wall formation in PX (Schneider et al., 2020). It is possible that ICR5 functions during this microtubule destabilization. It could be that the VND6-induced dedifferentiating cell culture system that has been used in previous studies (Oda and Fukuda, 2012) does not fully recapitulate the differentiating MX in the root and that these cells harbor characteristics of PX, which would have made it difficult to identify cell-specific functions of ICR5.

### The function of ICR2 in root hair growth

The split root hair phenotype of *icr2* mutant is not associated with changes in root hair density or position in the trichoblasts. This indicates that ICR2 function is required for maintenance of polar root hair elongation. The localization of ICR2 on microtubules in growing root hairs and the altered microtubule dynamics of *icr2* mutants (i.e., slower microtubule growth rate and increased rate of transitions between filament extension pause and shrinkage) indicate that ICR2 is necessary for stability of microtubules in root hairs. A split root hair phenotype has been associated with perturbation of microtubules and was described for several MAP mutants (Kang et al., 2017; Sakai et al., 2008; Whittington et al., 2001; Yang et al., 2007; Zhang et al., 2015). In root hairs, ICR2 on was found to localize on microtubules along the shank and not in active ROP domains at the root hair tip characterized previously (Jones et al., 2002).

Furthermore, although ICR2 was not recruited to active ROP2 domains, ROP2 gain-of-function mutations lead to formation of split root hairs (Jones et al., 2002; Kang et al., 2017), similar to the *icr2* mutant phenotype. Hence, ICR2 may function by recruiting active ROP2 from the plasma membrane to microtubules (Jones et al., 2002). Additionally, ROP10 was recently shown to regulate secondary cell wall formation in the shank leading to root hair shank hardening (Hirano et al., 2018). As we showed that ICR2 is recruited to active ROP10 domains, it is possible that ICR2 functions as a ROP10 effector in root hair. A recent study implicated ICR2 in recruitment of AGC1.5 where it phosphorylates ROPGEF4 and ROPGEF10 to promote root hair growth (Li et al., 2020). However, the split root hair phenotype of the *icr2* mutants, the localization of ICR2 on microtubules in root hairs, and the inability of active ROP2 domains to recruit ICR2 are not compatible with the proposed function of ICR2 in the activation of ROP2 function via AGC1.5 and ROPGEF4/10. Importantly, the distribution of ICR2 reported by Li *et al.* (Li et al., 2020) was determined by analysis of ICR2 with GFP fused to the N-terminal end; this fusion likely disrupted the interaction with microtubules, which takes place via the N-terminal end of ICR2. Hence, although the interaction of ICR2 with AGC1.5 is intriguing, its functional role will require additional investigation.

The analysis presented here together with previous works suggest that the ICR family proteins have multiple unique roles as ROP effectors, ROP regulators, and as MAPs that regulate microtubule dynamics. ICR2 and ICR5 may regulate microtubule destabilization through their interactions with proteins such as Kinesin13A. On the other hand, the analysis of microtubule dynamics and pit formation indicate that ICR2, and likely ICR3 and ICR5, stabilize microtubules and restrict ROP-mediated signaling by moving ROPs from the plasma membrane to microtubules. The tissue- and cell-specific functions of ICR2 and ICR5 may reflect interactions with different proteins in different cells. In addition, that only a subset of ROPs recruit microtubule-associated ICR2 to ROP domains may also affect the signaling specificity. Previously, we showed that ICR1 and ICR2 do not interact with the same proteins (Lavy et al., 2007), supporting the hypothesis that they function as differential adaptors of ROP signaling. The differences in the binding affinity of ICR1 and ICR2 to microtubules indicate that, in addition to interactions with different proteins, the two ICRs affect microtubules differently and may have different distributions in the cell. Although ICR2 localized to microtubules in all stages of the cell cycle, we did not detect any cell division abnormalities in the *icr2* single mutants or in the *icr2*/*icr5* double or *icr2*/*icr5*/*icr3* triple mutants. The function of ICR2 during cell division may be redundant with proteins outside the ICR family or may be required under specific conditions; this will be the focus of future studies.

## Materials and methods

### Molecular procedures

#### Plasmid DNA purification

Plasmid purification was carried out with a DNA-spin^TM^ Plasmid DNA Purification Kit (iNtRON biotechnology) according to the manufacturer’s protocol.

#### Polymerase chain reaction (PCR)

PCR was used for gene detection and cloning. For general uses such as colony screening, Taq DNA polymerase (Fermentas) was used. To eliminate error, for cloning purposes, PCR reactions were carried with the proof-reading Phusion DNA polymerase (NEB). Reaction conditions were according to the enzyme manufacturer's instructions with annealing temperatures chosen based on primers.

#### DNA fragment extraction from agarose gel

DNA extraction from agarose gels was done using the Gel extraction kit Qiaex II (Qiagen).

#### Cloning for co-expression, yeast two-hybrid, and BiFC assays

For imaging, *ROP6*, *ROP9*, *ROP10*, *ROP11* were subcloned downstream of *GFP* into *pGFP-MRC*. The resulting cassettes, containing a cauliflower mosaic virus (CaMV) *35S* promoter, the gene of interest, and a *NOS* transcriptional terminator, were subcloned into *pCAMBIA 2300* expression vector. Sequences encoding GFP-ROP2, GFP-ROP4, His-GAP1, His-GEF3_PRONE_, and ICR2-mCherry were cloned into expression vectors *pB7m34GW* or *pK7m34GW* by the Three-Way Gateway standard protocol (Invitrogen). The expression cassette included the *35S* promoter, the tag, the gene, and the NOS terminator. To obtain constitutively active ROP mutants, respective genes were mutated by site-directed mutagenesis. ROP6 and ROP10 were mutated to Q67L, ROP9 and ROP11 were mutated to G15V. For yeast two-hybrid analysis, ROP2, ROP4, ROP6, ROP9, ROP10, and ROP11 coding sequences were subcloned into *pGBT9.BS* and the ICR2 coding sequence was cloned into *pGAD GH*. For BiFC assays, YN-ROP2, YN-ROP4, YN-ROP6, YN-ROP9, YN-ROP10, YN-ROP11, ICR2-YC sequences were subcloned into pB7m34GW by the Three-Way Gateway standard protocol (Invitrogen). The expression cassette included the CaMV *35S* promoter, tag, gene of interest, and *NOS* terminator.

#### Creation of *pB7-pICR2::ICR2*

Intermediate vectors were created using Gateway BP Clonase. A 2493-bp fragment harboring the entire genomic sequence of *ICR2* (AT2G37080) from the ATG initiation codon through the stop codon was amplified from genomic DNA and subcloned into pDONR221. The promoter sequence of *ICR2* (2225 bp upstream of the *ICR2* initiation codon) was likewise amplified and subcloned into *pDONR-P4R1*. The *NOS* terminator was subcloned into *pDONR-P2R3*. All three intermediate vectors were further cloned into *pB7m34GW* destination vector using the Gateway LR Clonase II Plus Enzyme Mix for MultiSite LR recombination reaction.

#### Creation of *pK7-pICR2::ICR2-3xYPet* (EYFP variant)

Intermediate vectors were created using Gateway BP Clonase. A 2490-bp fragment harboring the entire *ICR2* (AT2G37080) genomic sequence from the ATG initiation codon but without the final stop codon was amplified from genomic DNA and subcloned into *pDONR221*. The promoter sequence of *ICR2* (2225 bp upstream of the *ICR2* initiation codon) was likewise amplified and subcloned into *pDONR-P4R*. 3xYPet-3xHA (Marquès-Bueno et al., 2016) was received from NASC (NASC code N2106295). This vector contains a 33-amino acid linker (DPAFLYKVARLEEFGTPGSKSISLDPLPAAAAA) between ICR2 and the three repeats of the fluorescent protein YPet to reduce potential steric hindrance. All three intermediate vectors were further cloned into *pK7m34GW* using the Gateway LR Clonase II Plus Enzyme Mix for MultiSite LR recombination reaction.

#### Creation of ICR1-His_6_ and ICR2-His_6_

*pET28b-ICR1-His_6_* was created by amplifying a 1050-bp fragment of the coding sequence of *ICR1* (AT1G17140) without a terminating stop codon from cDNA with flanking Nco1 and Not1 sites, and was subcloned into pJET1.2 (Thermo Scientific). The plasmid was than digested with Nco1 and Not1, and the *ICR1* fragment was subcloned into *pET28b*. *pET28b-ICR2-His_6_* was created by amplifying a 1749-bp fragment of the coding sequence of *ICR2* without the stop codon and subcloning into *pJET1.2*. It was than integrated into *pET28b* using primers containing overlapping sequence to *pET28b* and amplifying the entire vector by Transfer PCR (T-PCR) (Erijman et al., 2014).

#### Multiplex genome editing design and constructs

The polycistronic tRNA-gRNA system (PTG) was used to generate multiple sgRNAs with different target sequences by flanking the sgRNAs with a tRNA precursor sequence as previously described (Xie et al., 2015). A pJET-gRNA-tRNA plasmid, which contains a gRNA-tRNA-fused fragment, was used as a template to synthesize the PTG construct. The gRNA scaffold fragment was amplified by PCR using a pair of specific primers (Bsa-gRNA-F and gRNA-R), whereas the tRNA_Gly_ fragment was amplified as an overlapping fragment of the primers g-tRNA-F and tRNA-R. Then these two fragments were fused as a gRNA-tRNA by overlapping extension PCR using primers Bsa-gRNA-F and tRNA-R. The overlapping PCR product was separated and purified from an agarose gel, and then inserted into *pJET1.2* (Thermo Scientific) to generate the template plasmid. The specific spacer sequences targeting *ICR2*, *ICR3*, and *ICR5* were selected using the CRISPR-PLANT database (www.genome.arizona.edu/crispr/). The PTG clones were created using Golden Gate (GG) for the assembly of DNA fragments. In order to ligate multiple DNA fragments in a desired order, GG assembly requires distinct 4-bp overhangs to ligate two DNA fragments after digestion with BsaI. The gRNA spacers are the only unique sequences in the PTG and were used for this purpose. Each part was amplified with spacer-specific primers containing the BsaI adaptor, except two terminal parts using gRNA spacer primer and terminal specific primers containing BbsI site. These PCR fragments were ligated together using GG assembly to produce the PTG with complete gRNA spacers targeting *ICR2*, *ICR3*, and *ICR5*. The assembled product was amplified with short terminus specific primers containing the BbsI adaptor. Next, using a second GG assembly step, the PTG fragment was inserted into the BbsI digested *pEntr_L1L2_AtU6gRNA*. The PTG cassette was than inserted into *pMR294_pKGCAS9PLUS-1* by Gateway LR Clonase (Thermo Fisher Scientific). The *pEntr_L1L2_AtU6gRNA* and *pMR294_pKGCAS9PLUS-1* vectors were gifts from Professor Gitta Coaker, University of California, Davis.

In all cloning, PCR-generated fragments were sequenced to verify that no PCR-generated errors were introduced. In the cases of gene fusions, following cloning, the borders between fragments were sequenced to verify that fragments were in frame. All primers and plasmids used and generated in this work are listed in Tables S4-S6.

#### Sequencing

DNA sequencing was performed at the Tel Aviv University DNA sequencing facility and was carried using the BigDye Terminator Cycle Sequencing Kit (Applied Biosystems).

### Plant genomic DNA isolation

Typically, 100 mg of liquid N_2_ batch-frozen leaf tissue were ground with a mortar and pestle, and genomic DNA was isolated using the GenElute Plant Gemomic DNA Kit (Sigma) according to the manufacturer’s protocol.

### Total RNA isolation from plants

*Arabidopsis* seedlings were batch frozen using liquid N_2_, and tissue was ground with a mortar and pestle. Total RNA was isolated from the ground material using the RNeasy SV total RNA isolation kit (QIAGEN) according to the manufacturer’s instructions.

### RT-PCR (cDNA synthesis)

cDNA synthesis for standard RT-PCR experiments was carried out using High Capacity cDNA Reverse Transcription Kit (Applied Biosystems): 1 μg RNA dissolved in 10 μl H_2_O was added to 10 μl of the 2X Reverse Transcription Master Mix, containing 2 μl 10X RT buffer, 0.8 μl 25X dNTPs mix, 2 μl 10X RT Random Primers, 1 μl MultiScribe™ Reverse Transcriptase, 1 μl RNase Inhibitor, and 3.2 μl Nuclease-free H_2_O. The reaction was performed in a thermal cycler for 10 min at 25 °C, then 120 min at 37 °C, and 5 min at 85 °C (for inactivation).

### Sequence analysis

Sequence analysis was carried out using the SnapGene® (GSL Biotech; available at snapgene.com) sequence analysis software package. The BLAST algorithm (http://www.ncbi.nlm.nih.gov/BLAST) was used to search the DNA and protein database for similarity. Multiple sequence analysis was done using JALVIEW (Waterhouse et al., 2009) with the Clustal (Thompson et al., 1994) algorithm.

### Bacterial strains and growth conditions

*E. coli* DH5α(F’)-*F’* was used for heat shock transformation and molecular cloning. *Agrobacterium tumefaciens* strain *GV3101/pMP90* was used for transient and stable expression of recombinant genes in *N. benthamiana* and *Arabidopsis* as previously described (Lavy et al., 2007). Growth media for bacteria was prepared as previously described (Ausubel et al., 1995). For solid media, 1.5% w/v of agar was added to the medium. *E. coli* cells were selected on 100 μg/mL ampicillin or 50 μg/mL kanamycin. *Agrobacterium tumefaciens GV3101/pMP90* was selected on 100 μg/mL gentamycin and 50 μg/mL spectinomycin.

### Yeast two-hybrid assays

*Saccharomyces cerevisiae* strain *PJ69-4a* was used as host. Plasmids for expression of ROPS (*pGBT ROPs*) were co-transformed with *pGAD-ICR2* into yeast cells via a standard lithium acetate transformation protocol. Four decimal dilutions of colonies expressing both plasmids were grown on a medium lacking leucine (L), tryptophan (T), and histidine (H) supplemented with 1 mM 3-amino-1,2,4-triazole (3AT) for interaction detection or on −LT for growth monitoring. The plates were incubated at 28 °C.

### Plant materials and transformation

#### Plant materials

*Arabidopsis Col-0* ecotype was used as wild type in all experiments and was used for all transformation for generation of transgenic plants. *N. benthamiana* was used for transient expression in leaf epidermal cells. The *icr2-1(GK567F02), icr2-2(GK281B01)*, and *icr2-3(GK159B08)* T-DNA mutants were obtained from NASC and are in the *Col-0* background. In generating the CRISPR/Cas9-mediated genome edited mutants, transgenic plants were created by expression of appropriate gRNAs in a single transcriptional unit, spaced by tRNAs under the control of the *AtU6* promoter as described (Xie et al., 2015). The Cas9 in this system was expressed under the control of the *GEX1* egg-specific promoter, and therefore the genomic editing events identified were heritable and not somatic, thus improving the screening process. For analysis, the T-DNA containing the *pGEX1::Cas9-AtU6::tRNA-gRNA* expression cassette was crossed out from all mutants. Seeds for *UBN::RFP-MBD* were a gift from Dr. Sabine Müller, University of Tübingen and were previously described (Lipka et al., 2014). Plants used and generated in this work are listed in Table S7.

#### Plant growth conditions

Seeds of wild-type *Col-0*, transgenic, and mutant *Arabidopsis* plants were sown on the soil (Weizmann Institute mix, Pele Shaham, Ashkelon) and moved to stratification at 4 °C for 48 h in the dark in order to increase the uniformity of germination. The seeds were then moved to a growth chamber under long-day conditions (16 h light/8 h dark, light intensity 100 μE·m^−2^·s^−1^) at ~22 °C. *N. benthamiana* plants were grown in 10-cm pots. Seeds were sown on a mixture of 70% soil with vermiculite (Avi Saddeh mix, Pecka Hipper Gan). Plants were grown in an environmental growth chamber under conditions of long days (16 h light/8 h dark-light intensity 100 μE·m^−2^·s^−1^) at ~25 °C. For growth of *Arabidopsis* on plates, plates contained 0.5X Murashige Skoog (MS) medium (Duchefa) titrated to pH 5.7 with MES and KOH and 0.8% plant agar (Duchefa). In some cases, the medium was prepared with 1% sucrose. The seeds were then moved to a growth chamber and placed vertically in most cases, or horizontally for germination assays to grow under long-day conditions (16 h light/8 h dark-light intensity 100 μE·m^−2^·s^−1^) at ~22 °C. Prior to the transfer to the growth chamber, the sown seeds were stratified at 4 °C for 48 h in darkness. In both cases, seeds were surface sterilized by evaporation of HCl (6 mL) in sodium hypochlorite (100 mL) in a closed container for 1 h.

#### Stable transformation in *Arabidopsis*

Transformation was performed by the floral dip method (Clough and Bent, 1998).

#### Transient expression in *N. benthamiana*

Transient expression was carried out as previously described (Lavy et al., 2007).

### Vascular Cell Induction Culture System Using Arabidopsis Leaves (VISUAL)

Vascular cell dedifferentiation was carried out with the VISUAL system as previously described (Kondo et al., 2016) with the following modifications. In brief, 10-15 seedlings were grown in 10 ml liquid growth media and (2.2 g/L MS Basal Medium, 10 g/L sucrose and 0.5 g L^−1^ MES adjusted to pH to 5.7 using KOH) in 6-wells-plates at 25 ºC under a long-day regime (16 h light, 8 h dark) for 6 days until true leaves appeared. The central part of the hypocotyl was cut, and the roots were removed. For VISUAL induction, the aerial parts of the seedlings were incubated in induction medium (2.2 g/L MS Basal Medium containing 50 g/L D(+)-glucose adjusted to pH 5.7 with KOH) supplemented with 1.2 mg/L 2,4D, 0.25 mg/L kinetin and 10 μM bikinin in 12-well-plates for 4 additional days. Differentiating cells were imaged for lignin autofluorescence by excitation at 405 nm. Emission was detected with a spectral detector set between 410 nm and 524 nm. For simultaneous detection of lignin autofluorescence, chloroplasts, and ICR2-3xYPet, lambda mode was used with excitation at 514 nm. Emission was detected with a spectral detector set between 520 nm and 690 nm. Z-stacks were taken, and maximum intensity images were created.

### Analysis of microtubule dynamics

Microtubule dynamics was analyzed by high frequency time-lapse imaging of seedlings of *RFP-MBD*, *icr2-1xRFP-MBD*, and *icr2-2xRFP-MBD* genotypes at 8 DAG. Seedlings were grown on CellView^TM^ cell culture dishes, 35/10 mm, glass bottom (Grainer 627860) at a 45º angle, so that roots grew along the glass bottom between the growth medium and the glass. Imaging of root hairs and adjacent root epidermis cells was done at 2-s intervals, 60 frames total, using LSM 780-NLO confocal laser scanning microscope (Zeiss) in fast scanning mode with a 63X water immersion objective, and were visualized by excitation with an argon laser at 561 nm and spectral GaAsP detector set between 570 nm and 695 nm. Quantification of microtubule dynamics was done by tracking of individual microtubule filaments. The tracking data were used to create kymographs, which were then used to calculate microtubule growth and shrinkage rates, the time spent at each condition, as well as the transitions between them and pauses in growth/shrinkage. This analysis of imaging data was performed using the KymoToolBox ImageJ plugin (Zala et al., 2013). Typically, 5-10 microtubule filaments were analyzed per cell and five cells, each from a different plant, were analyzed for each genotype and cell type. Overall, the number microtubule filaments analyzed was 77-113.

### Secondary cell wall pits area and pit density per area

Analysis of secondary cell wall of the MX pits was carried out on seedling roots at 8 DAG. Roots were imaged using differential interference contrast (DIC) light microscopy after clearing with chloral hydrate:lactic acid (2:1) for 1-3 days. To quantify the area of secondary cell wall pits, pits were manually selected in DIC images and analyzed using ImageJ. Pit density was calculated as the number of secondary cell wall pits divided by the area of MX vessel cells and expressed as the number of pits per 1000 μm^2^. Two or three cells were imaged for each plant, and four or five plants were analyzed for each genotype.

### Protoxylem lignification

Roots at 7 DAG were imaged for lignin autofluorescence by excitation at 405 nm. Emission was detected with a spectral detector set between 410 nm and 524 nm. Z-stacks were taken of 6-10 focal planes, and maximum intensity images were created. Analysis was carried out in the maturation zone of the root on maturing PX cells, which at this region have a well-defined spiral pattern. No MX differentiation was detected. Mean distance between lignified spirals was measured using the semi-automated Cell-o-Tape macro for ImageJ (Fiji). Five roots were analyzed for each genotype, and in each plant two PX cells were imaged and quantified.

### Analysis of root hairs deformation

Root hairs were observed in seedlings grown on 0.5X MS agar medium at 8 DAG. Ten seedlings for each genotype were compared, and the percent of normal and deformed root hairs was scored.

### Root hair measurements

Root hairs in seedlings at 7 DAG were imaged and measured as previously described (Denninger et al., 2019). The first visible swelling of the cell outline was defined as first bulge, and distance to root tip was measured. Root hair density was analyzed in the next 2 mm. Root hair length was measured in a region 3-6 mm away from the root tip.

### Light and confocal laser scanning microscopy

Brightfield and Nomarsky DIC imaging was performed with an Axioplan-2 Imaging microscope (Zeiss) equipped with an Axio-Cam and a cooled charge-coupled device camera using either 10X, 20X dry, or 63X water immersion objectives with numerical aperture values of 0.5, 0.9, and 1.2, respectively. Laser scanning confocal microscopy and associated brightfield and DIC imaging was performed using LSM 780-NLO confocal laser scanning microscope (Zeiss) with 10X and 20X air objectives and 40X and 63X water immersion objectives with numerical apertures of 0.3, 0.8, 1.2, and 1.15. Fluorescein was visualized by excitation with an argon laser at 488 nm; emission was detected between 493 and 556 nm. Rhodamine was visualized by excitation with an argon laser at 561 nm; emission was detected between 566 and 685 nm. 3xYPet was visualized by excitation with an argon laser set at 514 nm; emission was detected 526 and 570 nm.

### Image analysis

Image analyses were performed with ZEN 2012 Digital Imaging, Photoshop CS5.1 (Adobe Systems) and ImageJ (FIJI).

### Quantifications and statistical analyses

Stacked charts and box plots were prepared using JMP (SAS) or Office Excel 2016 (Microsoft). Statistically significant differences were determined using the Student’s *t*-test or one-way ANOVA and Tukey-HSD post-hoc analysis, as noted in the figure legends and Tables S2 and S3.

### Expression of ICR1-His6 and ICR2-His6 in *E. coli*

ICR1-His_6_ and ICR2-His_6_ were transformed into the BL21 (Rosetta) *E.coli* strain. Cells were grown at 37 °C to an OD_600_ of 0.5-0.7 and then induced with 1 mM isopropyl-beta-D-thiogalactopyranoside overnight at 16 °C. Immediately after induction cells were harvested by centrifugation at 5000 X *g* for 15 min at 4 °C, and stored at −80 °C until further use.

### Purification of ICR1-His6 and ICR2-His6

Protein purification was carried out with the AKTA-prime protein purification system (GE Healthcare). First, cells were homogenized by sonication using VCX500 ultrasonic processor (Sonics & Materials, Inc.) in washing buffer (50 mM NaH_2_PO_4_, 300 mM NaCl, 20 mM imidazol, and 5% glycerol, pH 8.0) containing 1 mM DTT. ICR1-His_6_ and ICR2-His_6_ recombinant proteins were purified over a His-TRAP FF column (GE Healthcare) with 1-mL bed volume. The column was washed with 30 mL of washing buffer. The proteins were released with Imidazole Elution buffer (50 mM NaH_2_PO_4_, 300 mM NaCl, 250 mM imidazole and 5% glycerol, pH 8.0). The proteins were concentrated with Amicon Ultra-15 filters (Millipore) with molecular weight cutoffs of 30 kDa and 50 kDa for ICR1-His_6_ and ICR2-His_6_, respectively, at 4,000 *g* and 4 °C to a final volume of approximately 500 μl. The concentrated protein samples were filtrated through Millex 0.22-μm syringe filters (Millipore) and loaded onto a Superdex 200 HR 10/30 gel-filtration column (GE Healthcare) and eluted with a gel filtration column buffer (50 mM NaH_2_PO_4_, pH 7.0). To concentrate the protein, an Amicon Ultra-15 Centrifugal Filter Device was used, protein was centrifuged, and the buffer was exchanged to PEM buffer (0.1 M PIPES, 1 mM EGTA, and 1 mM MgCl_2_, pH 6.9). Purified proteins were again concentrated using the Amicon Ultra-15, divided into aliquots, batch frozen in liquid nitrogen, and kept at −80 °C until further use. Protein concentrations were determined using the BCA Protein Assay kit (Pierce) according to the manufacturer's protocol.

### Microtubule co-sedimentation assay

Porcine brain tubulin was purified as described (Castoldi and Popov, 2003). For the co-sedimentation assay, 0, 1, 2, 4, 6, 8, and 10 μM purified ICR1-His_6_ or ICR2-His_6_ were added to taxol-stabilized microtubules (5 μM tubulin) in PEMT (100 mM PIPES, 1 mM EGTA, 1 mM MgCl_2_, 1 mM GTP, and 20 μM taxol, pH 6.9). The samples were centrifuged at 100,000 *g* at 25 °C for 15 min. Pellets and supernatants were analyzed by 10% SDS-PAGE and visualized by staining the gels with Coomassie Brilliant Blue R 250. His-NtMAP65-1c and BSA were used as positive and negative controls, respectively.

### Microtubule immunofluorescence co-localization and *in vitro* bundling assays

Rhodamine-labeled tubulin was prepared as previously described (Hyman, 1991). For the co-localization assay, taxol-stabilized microtubules composed of tubulin mixed with rhodamine-labeled tubulin (molar ratio, 1:4) in PEMT were incubated with 3 μM ICR1-His_6_ or 0.5 μM ICR2-His_6_ for 15 min at 37 °C and then crosslinked with 20 mM 1-ethyl-3-(3-dimethylaminopropyl) carbodiimide (Pierce Biotechnology) for 5 min at 37 °C. The mixture was then centrifuged at 12,000 *g* for 5 min, and the pellet was resuspended in PEM buffer preheated to 37 °C. ICR1-His_6_ and ICR2-His_6_ were stained with an anti-His antibody (1:5,000) and secondary antibody conjugated with fluorescein (Sigma; F0257; 1:5,000). The solution was then centrifuged at 12,000 *g* for 5 min, and the pellet was resuspended with PEM buffer preheated to 37 °C. An aliquot of 1 μl was put on a poly-L-lysine coated glass slide (Sigma, P0425) and observed by confocal microscopy. For the *in vitro* bundling assay, the same taxol-stabilized rhodamine-labeled microtubules were incubated with 0.1, 0.5, 1, and 5 μM ICR1-His_6_ or 0.1, 0.5, 1, and 2 μM ICR2-His_6_ for 30 min at 37 °C and then treated with 0.005% glutaraldehyde. A 1-μl aliquot of each sample was put on a poly-L-lysine coated glass slide (Sigma, P0425) and observed by confocal microscopy.

## Acknowledgements

This research was supported by ISF-NSFC grant (3342/20 to SY and 32061143018YF and ISF 2144/20 to SY. We thank Professor Gitta Coaker, UC Davis, and Dr. Sabine Muller, University of Tuebingen for providing materials.

## Supplement figures

**Figure 1-Supplement 1:**
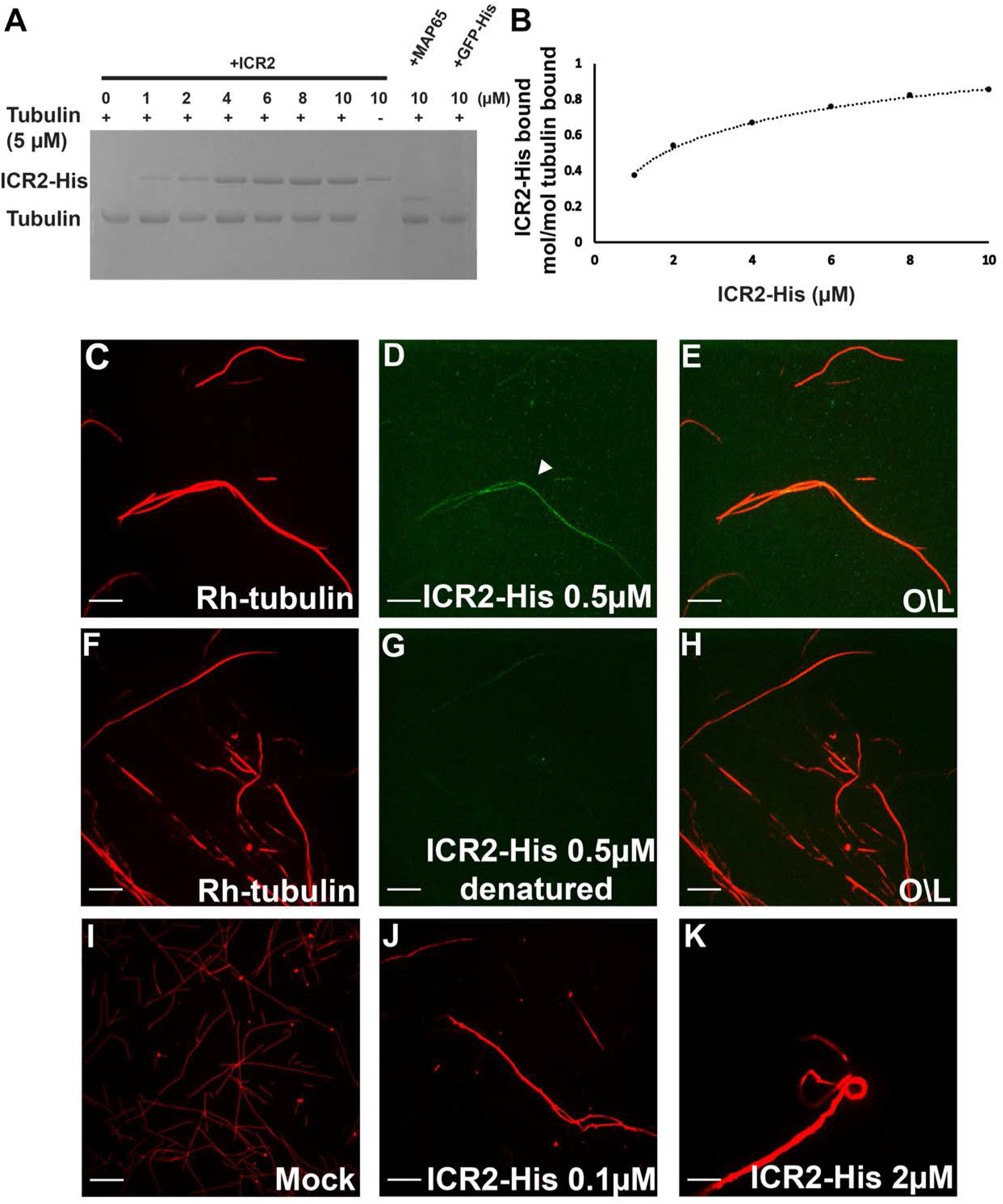
*ICR2* expression pattern and during development. Transcriptomics data of *ICR2* expression levels. Figure adapted from the *Arabidopsis* eFP Browser (https://bar.utoronto.ca/efp/cgi-bin/efpWeb.cgi).

**Figure 2-Supplement 1:**
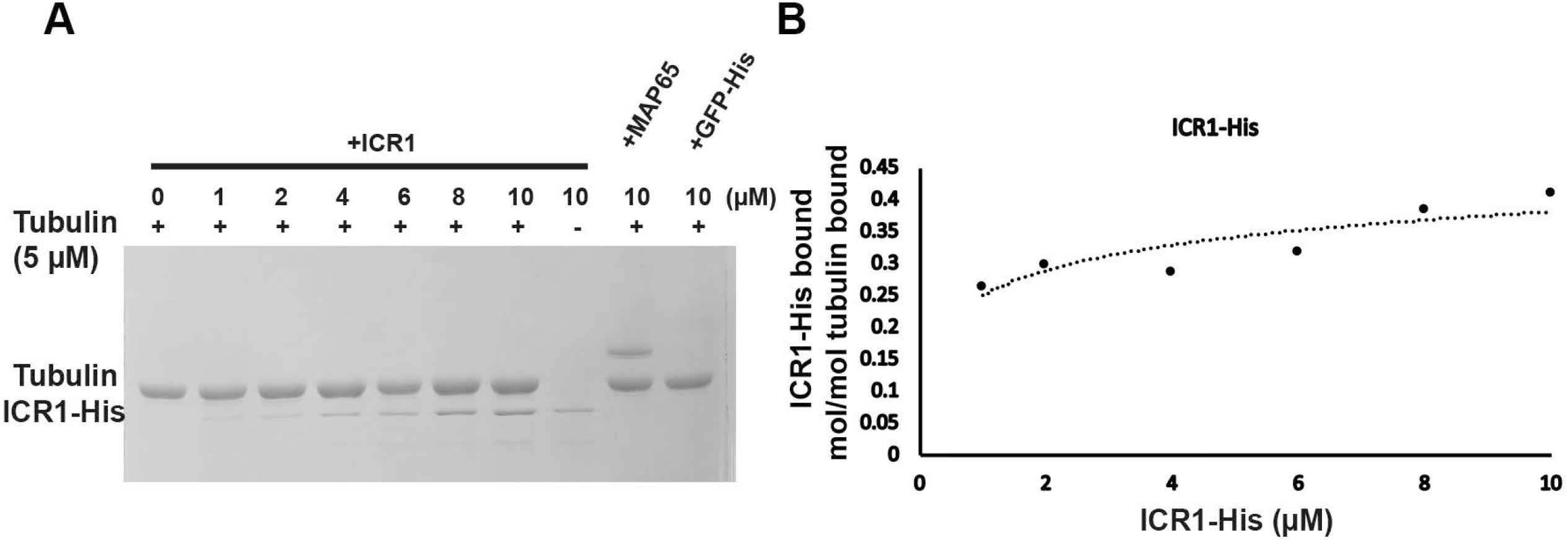
Map of the *ICR2* locus showing three *icr2* T-DNA insertion mutant alleles. **(A)** *icr2-1* (*GK567F02*) T-DNA insertion is in the third exon, 297 bp before the stop codon; *icr2-2* (*GK281B01*) insertion is in the first exon, 19 bp after the initiation codon; *icr2-3* (*GK159B08*) insertion is in the third exon, 472 bp after the start of the exon. **(B)** *icr2* mutant plants have no *ICR2* mRNA transcript (1,750 bp). *Ubiquitin 5* (UBQ) was detected as a control (650 bp).

**Figure 2-Supplement 2:**
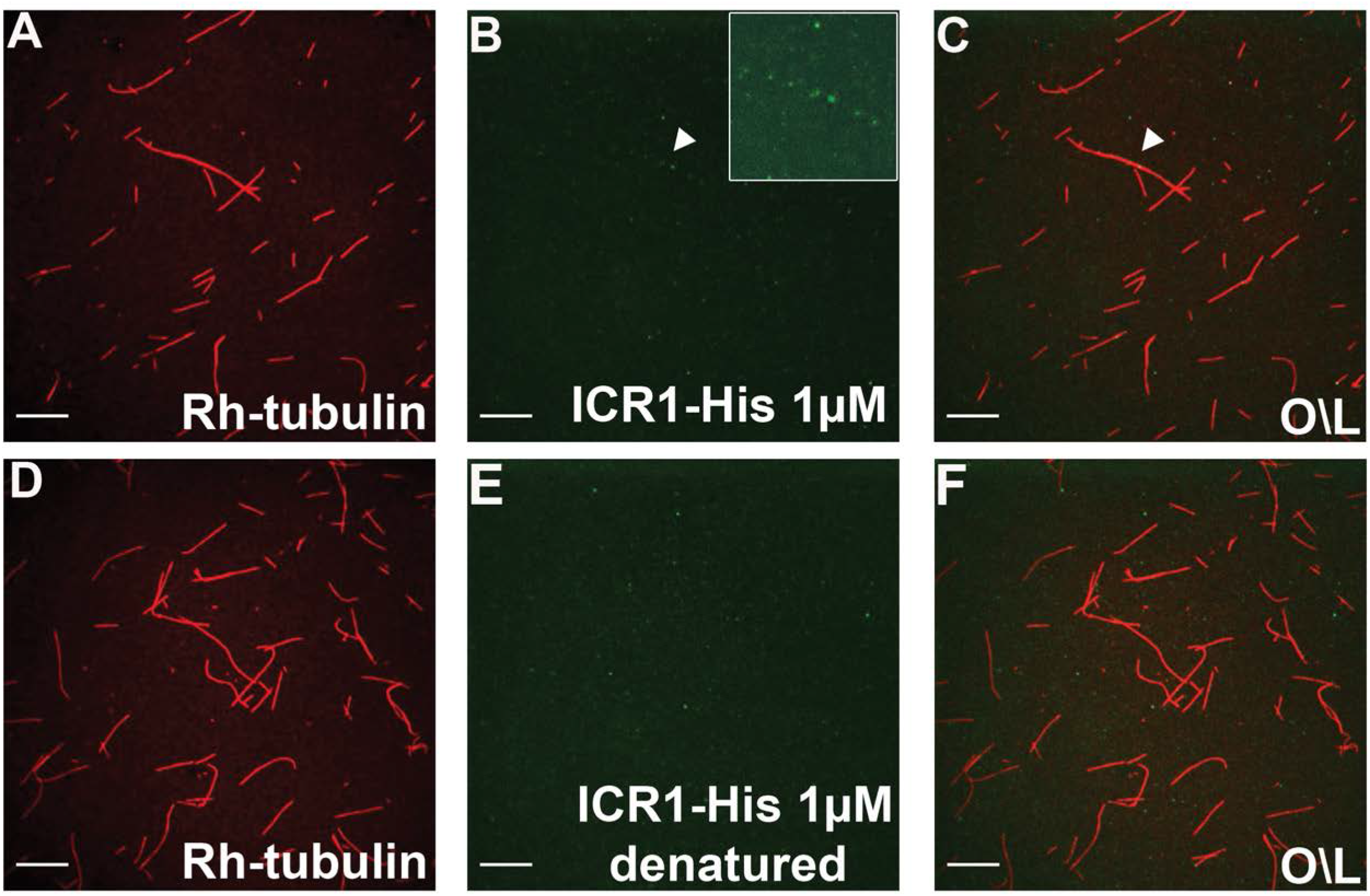
The CRISPR/Cas9 generated mutations in *ICR2*, *ICR3,* and *ICR5*. **(A)** Positions of the gRNA target sequence for each gene. (**B)** Sequences of InDels in the mutant alleles aligned with the WT *Col-0* allele. Inserted bases are marked in red. Dashed lines indicate deletions. **C)** Predicted amino acid sequences of the mutants. Asterisk indicates stop codon.

**Figure 4-Supplement 1:**
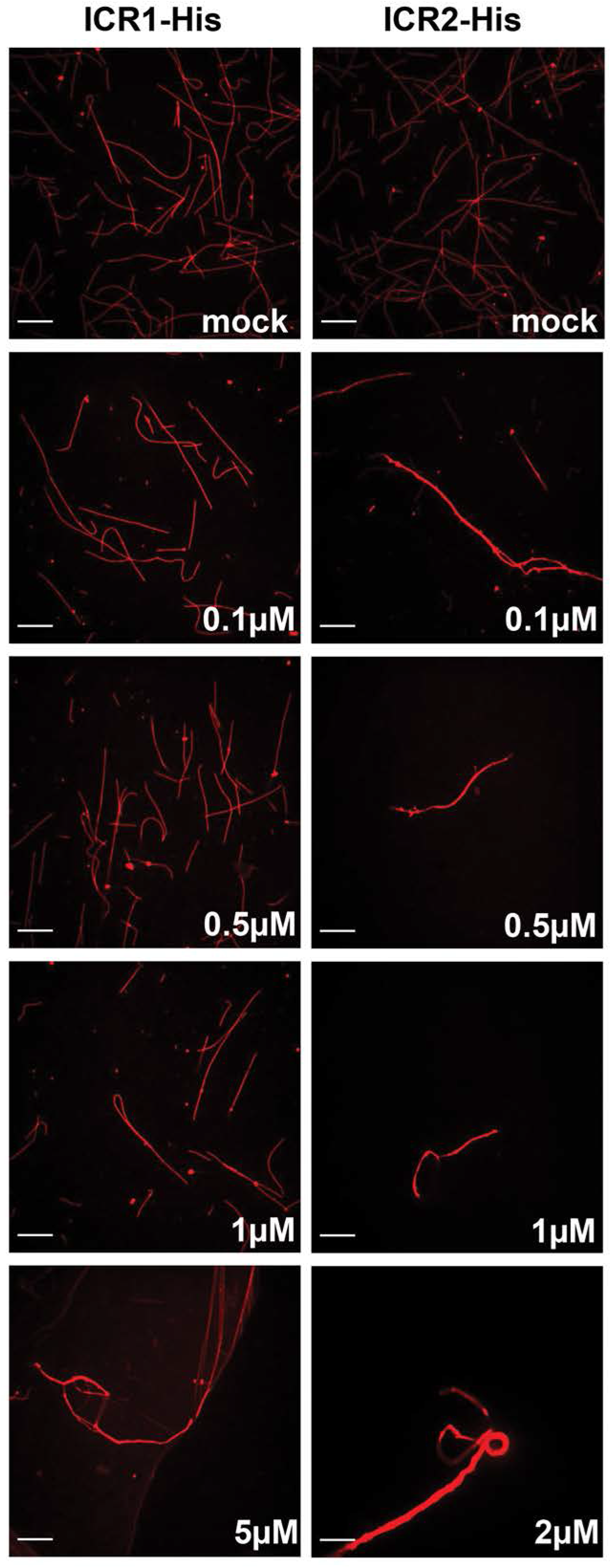
Root hair initiation sites, density, and length in WT and single, double, and triple mutants. Quantification of **(A)** normalized distance of first bulge from root tip (n≥16), **(B)** length of root hairs (n≥142), and **(C)** density of root hairs (n≥16). No significant differences were identified between the lines using ANOVA. The boxes are the interquartile ranges, the whiskers represent the 1^st^ and 4^th^ quartiles, and the lines are the averages. See source data-Figure 4-Supplement 1 A,B,C.

**Figure 5-Supplement 1:**
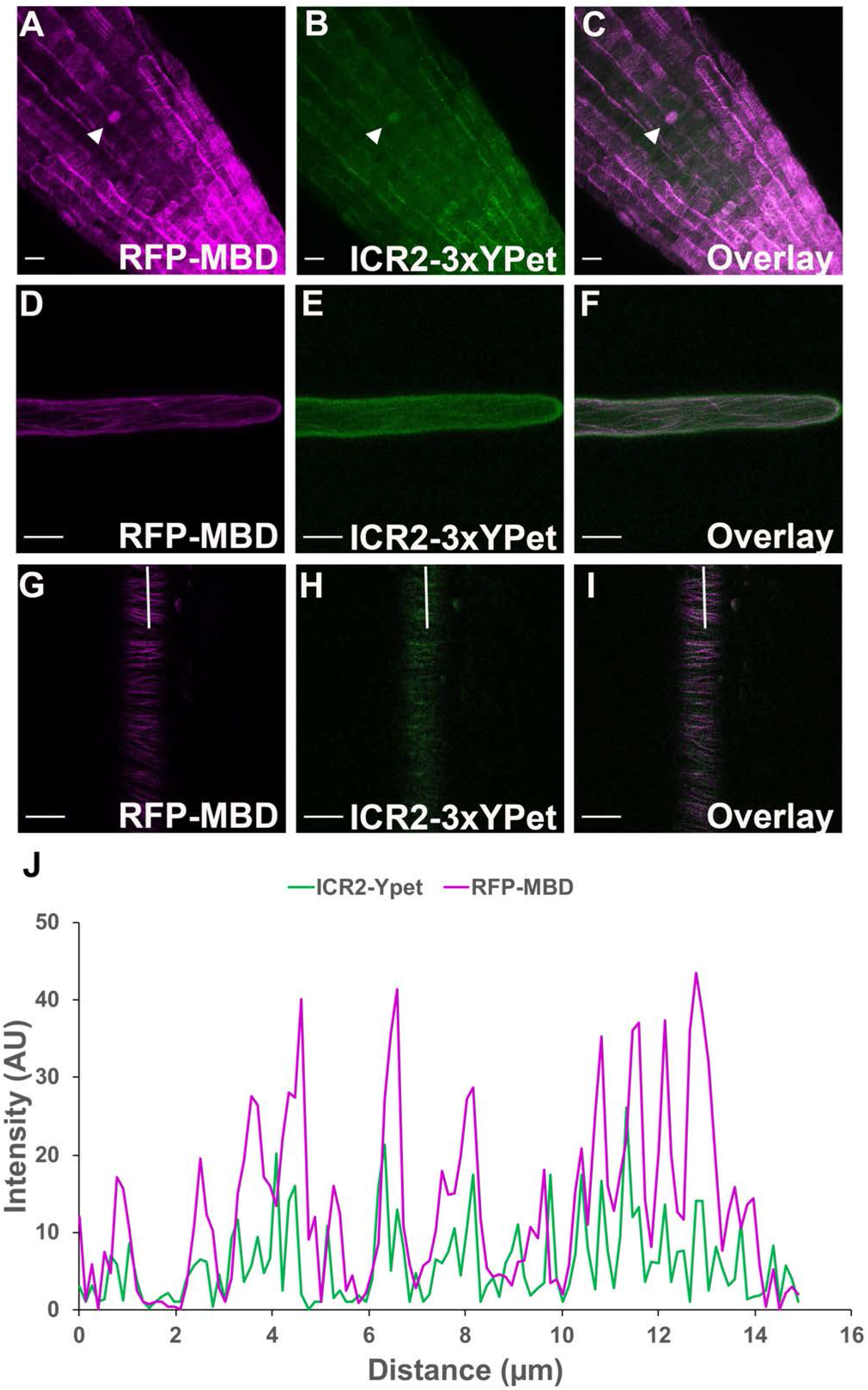
ICR2 localization with a microtubule marker is disrupted by oryzalin. **(A)** Image of *N. benthamiana* leaf epidermis that transiently expresses ICR2-3xYPet. **(B)** Image of *N. benthamiana* leaf epidermis that transiently expresses ICR2-3xYPet after oryzalin treatment, which disrupts microtubules. **(C-E)** Image of *N. benthamiana* leaf epidermis that transiently expresses ICR2-3xYPet and RFP-MBD. **(F-H)** Image of *N. benthamiana* leaf epidermis that transiently expresses ICR2-3xYPet and RFP-MBD after oryzalin treatment. Scale bars are 10 μm.

**Figure 7-Supplement 1:**
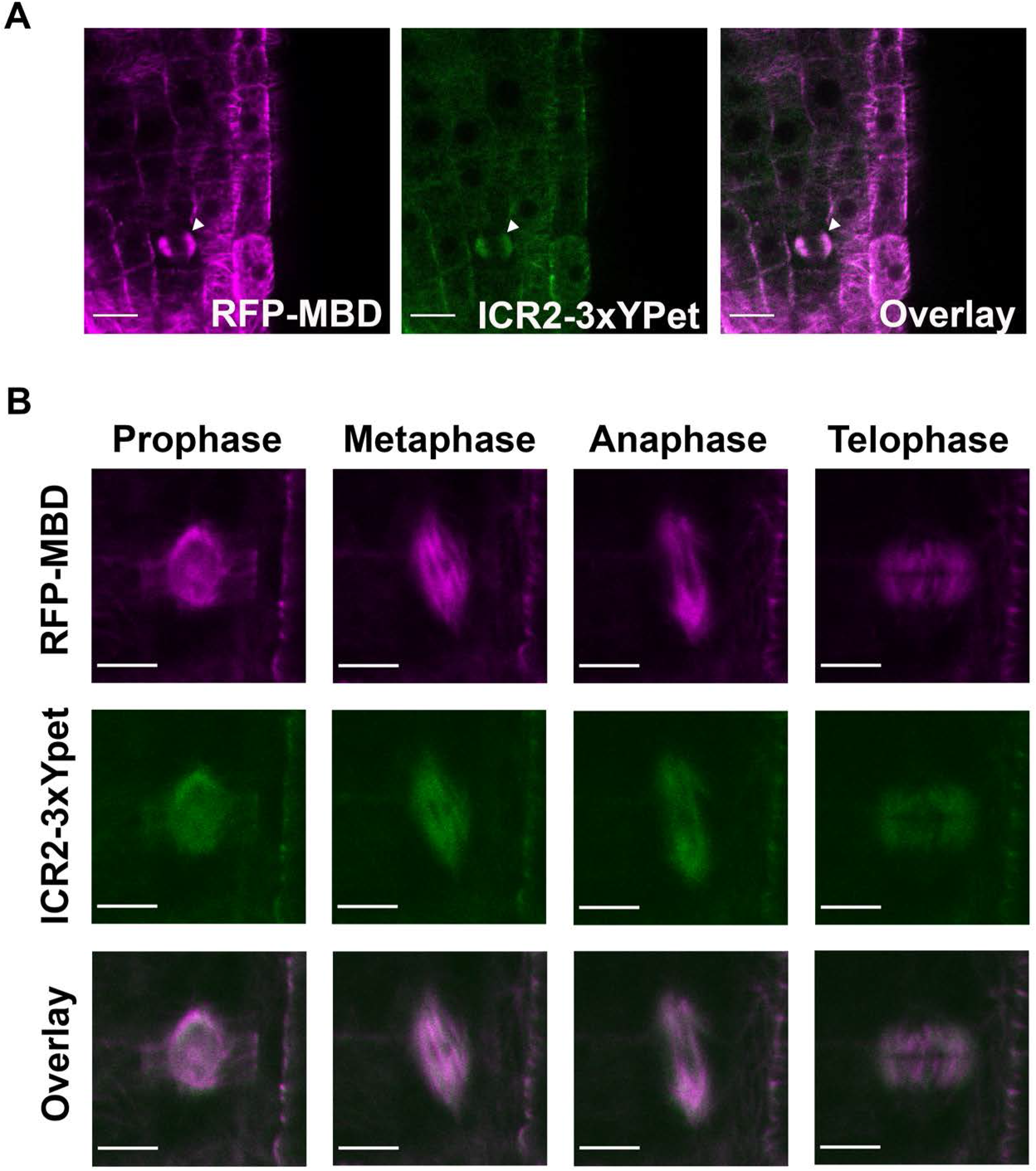
ICR1 interacts with microtubules *in vitro*. (**A**) Coomassie brilliant blue stained SDS-PAGE of recombinant ICR1-His_6_ co-sedimented with taxol-stabilized microtubules pre-polymerized from 5 μM tubulin. His-AtMAP65-1 was used as a positive control and GFP-His as a negative control. (**B**) Quantification of the ICR1-His_6_ band. The plot averages three replicates. See source data-Figure 7-Supplement 1 B.

**Figure 7-Supplement 2:**
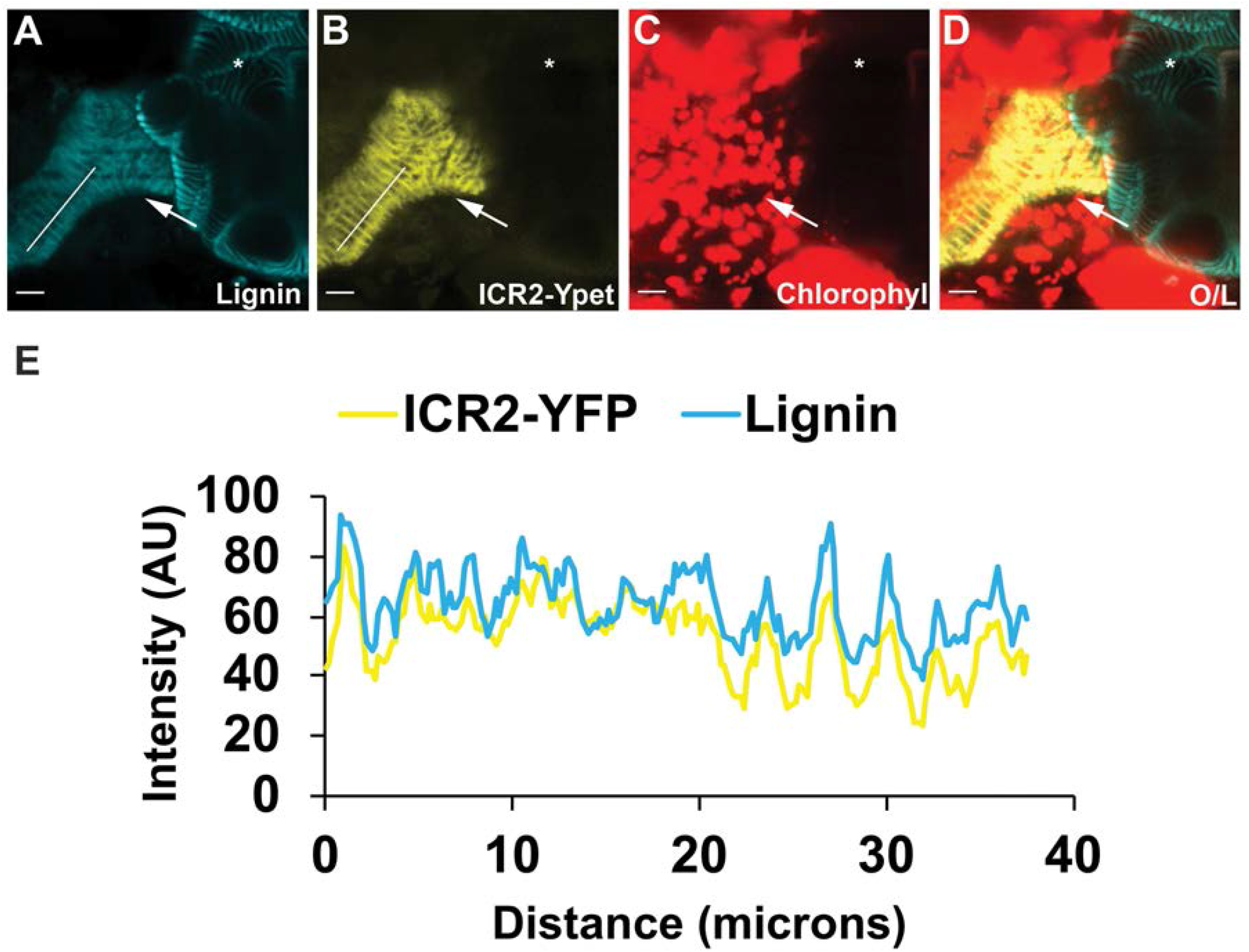
ICR1 co-localizes with microtubules *in vitro*. Immunofluorescence images showing the specific binding of (**A-C**) recombinant fluorescein-labeled ICR1-His_6_ (green) along rhodamine-labeled microtubules (red). Inset in panel B is an enlargement of the region highlighted by arrowheads in B and C. (**D-F**) Imaging of denatured ICR1-His used as control. Scale bars, 10 μm in all panels.

**Figure 7-Supplement 3:**
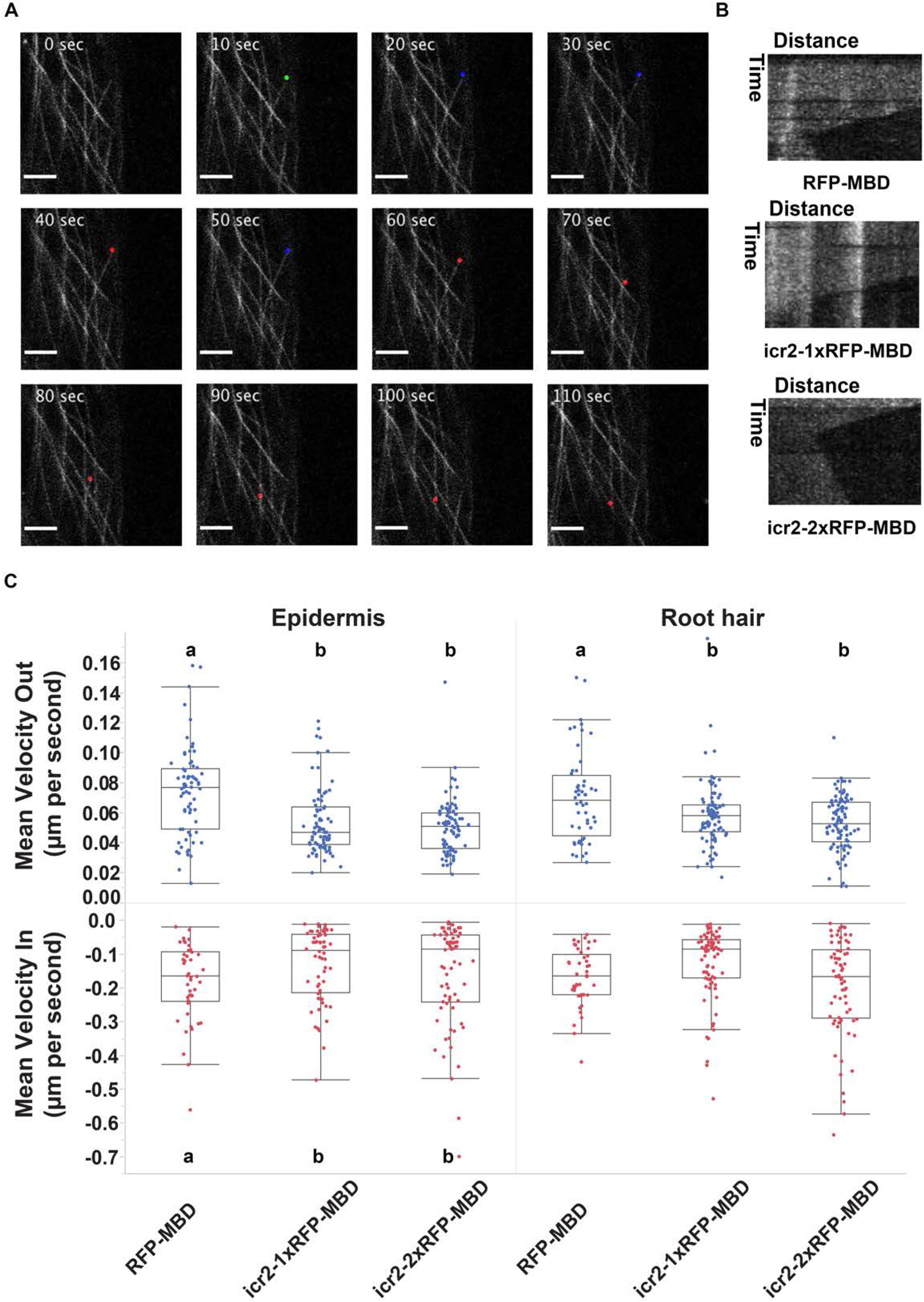
ICR1 and ICR2 induce microtubule bundling. Rhodamine-labeled microtubules (red) formed bundles when incubated with a range of concentrations of recombinant ICR1-His_6_ and ICR2-His_6_. The panels displaying mock, 0.1 μM ICR2, and 2 μM ICR2 samples are duplicates of Figure 7 panels I, J, and K, respectively, and are presented here for the sake of clarity.

**Figure 11-Supplement 1.**
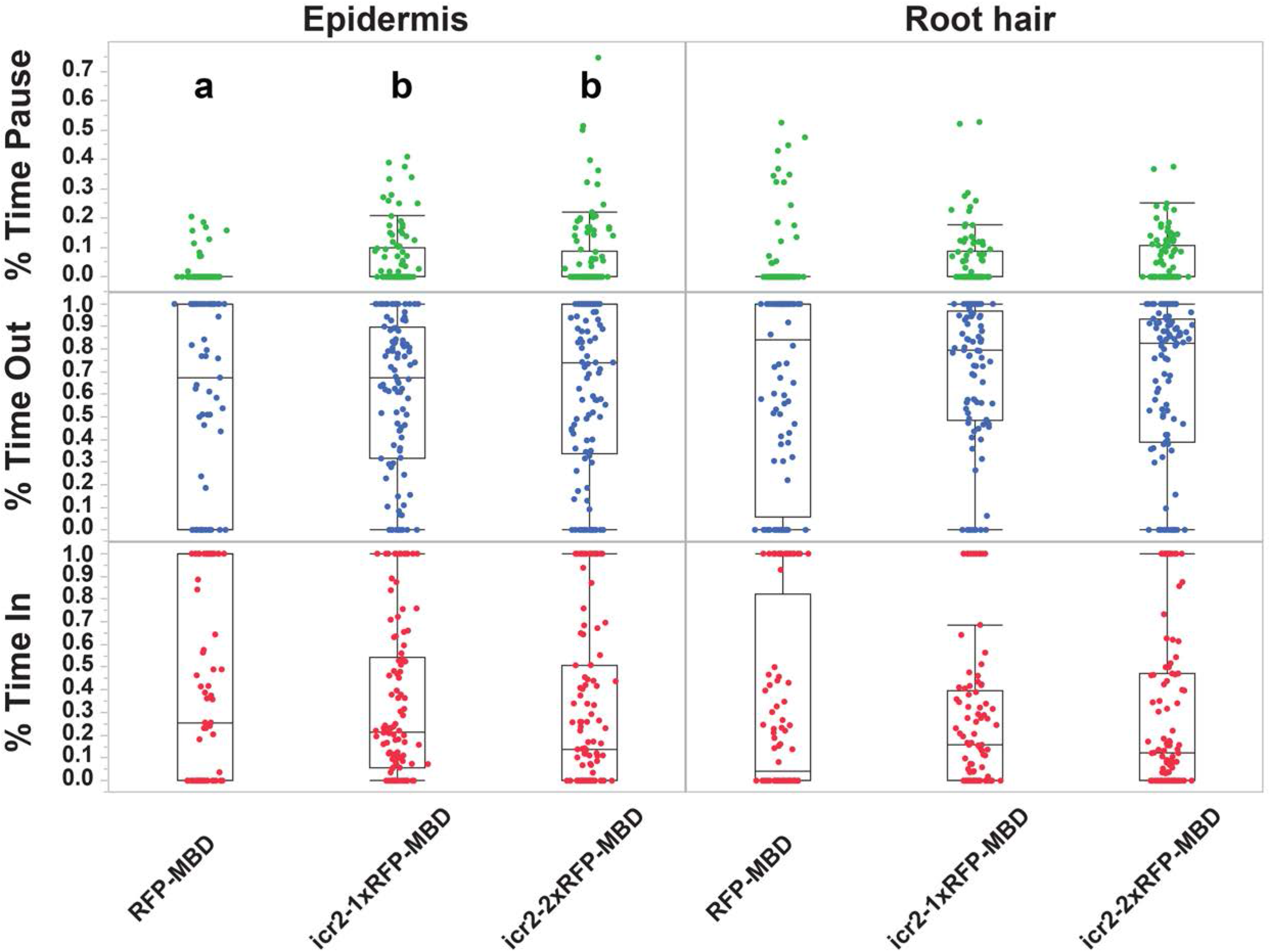
ICR2 affects pause time of microtubules between growth and shrinkage. Fraction of the time microtubule filaments were extending (blue), pausing (green) or shrinking (red). Means with different letters are significantly different (Tukey’s HSD, p<0.05). One-way ANOVA results are in Table S3. The boxes are the interquartile ranges, the whiskers represent the 1^st^ and 4^th^ quartiles, and the lines are the averages. n≥77 for each genotype. See source data-Figure 11-Supplement 1.

**Figure 11-Supplement-2:**
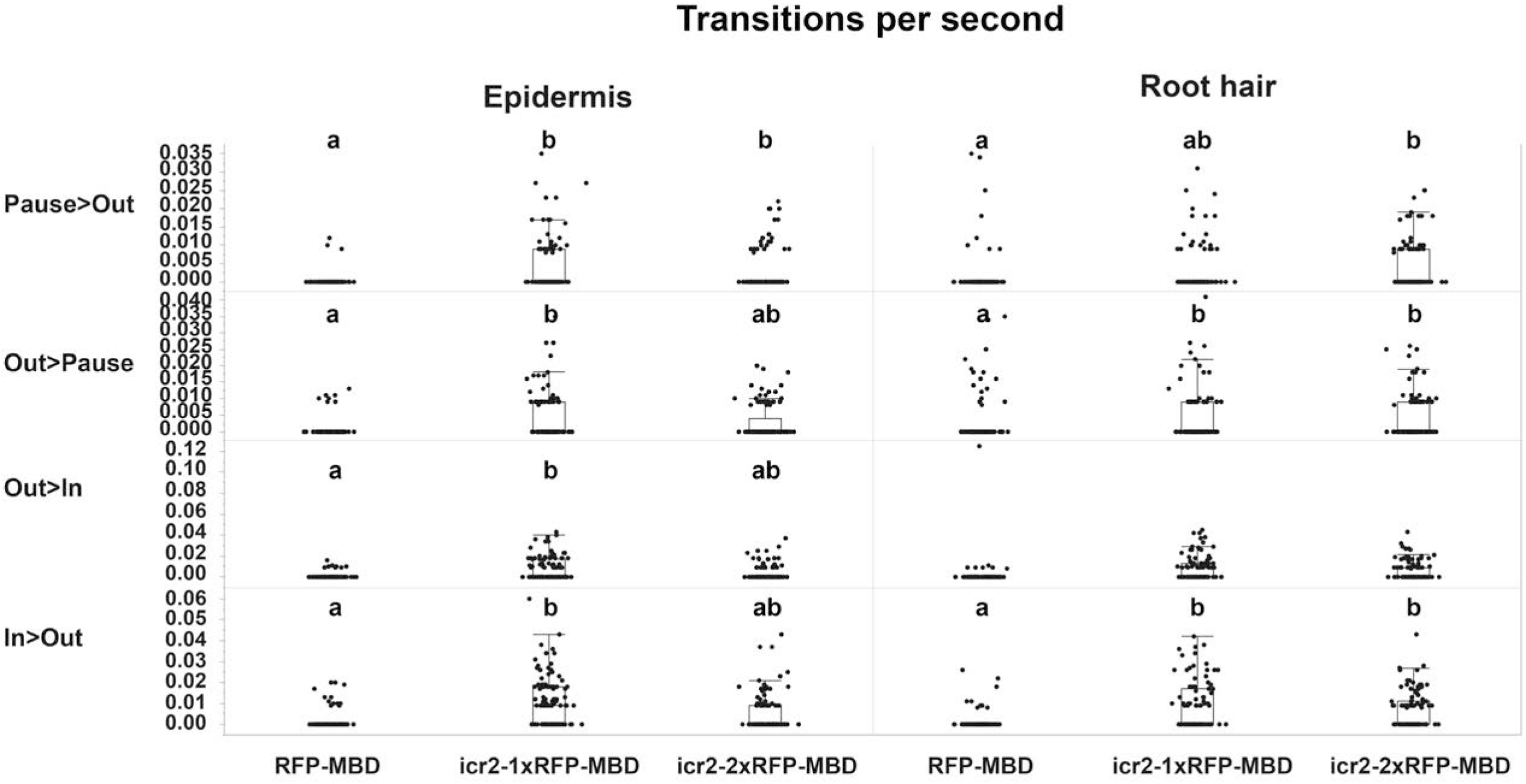
The effects of ICR2 on microtubule dynamics as detected by frequency of transitions. Number of transitions per second for each genotype at each cell type. In, shrinkage; Out, extension, Pause, pause. Means with different letters are significantly different (Tukey’s HSD, p<0.05). One-way ANOVA values are in Table S3. The boxes are the interquartile ranges, the whiskers represent the 1^st^ and 4^th^ quartiles, and the lines are the averages. n≥77 for each genotype. See source data-Figure 11-Supplement 2.

